# The effect of acquisition resolution on orientation decoding from V1 BOLD fMRI at 7 Tesla

**DOI:** 10.1101/081604

**Authors:** Ayan Sengupta, Renat Yakupov, Oliver Speck, Stefan Pollmann, Michael Hanke

**Author notes:** Corresponding author Email address (Ayan Sengupta).

## Abstract

A decade after it was shown that the orientation of visual grating stimuli can be decoded from human visual cortex activity by means of multivariate pattern classification of BOLD fMRI data, numerous studies have investigated which aspects of neuronal activity are reflected in BOLD response patterns and are accessible for decoding. However, it remains inconclusive what the effect of acquisition resolution on BOLD fMRI decoding analyses is. The present study is the first to provide empirical ultra high-field fMRI data recorded at four spatial resolutions (0.8 mm, 1.4 mm, 2 mm, and 3 mm isotropic voxel size) on this topic — in order to test hypotheses on the strength and spatial scale of orientation discriminating signals. We present detailed analysis, in line with predictions from previous simulation studies, about how the performance of orientation decoding varies with different acquisition resolutions. Moreover, we also examine different spatial filtering procedures and its effects on orientation decoding. Here we show that higher-resolution scans with subsequent down-sampling or low-pass filtering yield no benefit over scans natively recorded in the corresponding lower resolution regarding decoding accuracy. The orientation-related signal in the BOLD fMRI data is spatially broadband in nature, includes both high spatial frequency components, as well as large-scale biases previously proposed in the literature. Moreover, we found above chance-level contribution from large draining veins to orientation decoding. Acquired raw data were publicly released to facilitate further investigation.

## Introduction

The term multivariate pattern (MVP) analysis summarizes a range of data analysis strategies that are highly suitable for studying neural representations encoded in distributed patterns of brain activity (see, for example, Haxby, 2012; Haynes, 2009; Zhang et al., 2015; Bonte et al., 2014). While there is an ever increasing number of publications that demonstrate the power of MVP analysis for functional magnetic resonance imaging (fMRI) data (Op de Beeck, 2010; Freeman et al., 2011; Alink et al., 2013; Freeman et al., 2013) with standard resolution (a voxel size of about 2-3 mm isotropic), MVP analysis is especially promising in the context of high-resolution fMRI. Ongoing technological improvements, such as ultra high-field MRI scanners (7 Tesla or higher) have pushed the resolution for fMRI to a level that is approaching the spatial scale of the columnar organization of the brain (Yacoub et al., 2008; Heidemann et al., 2012). Being able to use fMRI to sample brain activity patterns at a near-columnar level makes it feasible to employ MVP analysis with the aim to decode distributed neural representations of an entire cortical field at a level of detail that is presently only accessible to invasive recording techniques with limited spatial coverage. However, at this point, it is un-clear which spatial resolution is most suitable for decoding neural representation from fMRI data with MVP analysis. While higher resolutions can improve the fidelity of the BOLD signal by, for example, reducing the partial volume effect (Weibull et al., 2008), the benefits can be counteracted by physiological noise (such as inevitable motion) and lower temporal signal-to-noise ratio (tSNR). This raises the question: does the decoding of neural representations continuously improve with increasing spatial resolution, or is there an optimal resolution for a given type of representation?

In this study, we aim to address this question for the most frequently employed MVP analysis technique: a cross-validated classification analysis, where a classifier is repeatedly trained to distinguish patterns of brain activation from fMRI data of a set of stimulus conditions, and its prediction accuracy is evaluated against a left-out data portion (Pereira et al., 2009). We selected oriented visual gratings in primary visual cortex as decoding subject, because it is likely to be the most extensively studied paradigm regarding the application of MVP analysis on fMRI data, starting with the classic studies of Kamitani and Tong (2005) and Haynes and Rees (2005). It was shown that orientation can be decoded reliably at resolutions ranging from standard 3 mm isotropic voxels in the aforementioned studies, to 1 mm (Swisher et al., 2010), and that it is possible to directly measure orientation columns in V1 with 7 Tesla fMRI of 0.5 × 0.5 mm (in-plane) resolution (Yacoub et al., 2008; Uğurbil, 2012). These findings led to a discussion on the origin and the spatial scale of the signals that classifiers can use to learn to discriminate different orientations (*e.g.*, Op de Beeck, 2010; Swisher et al., 2010; Alink et al., 2013; Freeman et al., 2013). To investigate these questions, the authors typically acquired high-resolution fMRI and simulated a lower-resolution acquisition by applying spatial filters to the original data (see Swisher et al., 2010), or reconstruction of k-space data to different resolutions (Gardumi et al., 2016), in order to compare metrics, such as prediction accuracy, across a range of spatial frequencies. However, these approaches have not gone unchallenged as it is unclear to what degree particular filtering strategies (*e.g.*, Gaussian filtering vs. low-pass filtering in the spatial frequency domain, see Misaki et al., 2013) can effectively simulate the properties of fMRI recorded at a lower physical resolution, where a change in slice thickness alone can significantly alter image contrast. Despite this criticism, we are not aware of any study that has compared the performance of orientation decoding in visual cortex across a range of physical acquisition resolutions.

In this study, we provide empirical data on the effect of spatial acquisition resolution on the decoding of visual orientation from high field (7 Tesla) fMRI. We recorded BOLD fMRI data at 0.8 mm, 1.4 mm, 2 mm and 3 mm voxel size while participants were visually stimulated with oriented phase-flickering gratings using a uniform event-related paradigm. Chaimow et al. (2011) investigated the effect of acquisition resolution on decoding of the stimulated eye using simulated 3 Tesla fMRI data based on a model of ocular dominance columns. They found that a resolution of 3 mm was optimal for decoding and performance decreased with higher or lower resolution. It is known that the organization of orientation columns is characterized by higher spatial frequencies than ocular dominance columns (Obermayer and Blasdel, 1993) and the BOLD point-spread function (PSF) is considerably smaller than that at 3 Tesla (≈ 2.3 mm FWHM vs. ≈ 3.5 mm FWHM Shmuel et al., 2007; Engel et al., 1997). Considering that, we expect the maximum orientation decoding accuracy to be observed at a resolution higher than 3 mm

The primary purpose of this study is to explore how spatial resolution as an acquisition parameter, or as a preprocessing outcome impacts decoding. These multiresolution data allow for evaluating filtering strategies used in previous studies in terms of their validity regarding the simulation of lower-resolution fMRI acquisitions from high-resolution data. These data also enable the investigation of the contributions of discriminating signal from individual spatial frequency bands for each resolution. Moreover, we collected high-resolution susceptibility weighted imaging data for blood-vessel localization in order to investigate the role of large draining veins that may carry orientation-discriminating signals reflected in low spatial frequency components when sampled by millimeter range voxels (Kamitani and Tong, 2005; Kriegeskorte and Bandettini, 2007; Shmuel et al., 2010; Gardner, 2010). In combination with the multi-resolution fMRI data, we can investigate the effect of this potential signal source on the orientation decoding at a range of of spatial scales.

While our primary focus is on the technical aspect of acquisition resolution for decoding information from BOLD signal patterns using the representation of visual orientations as a well-researched example, we acknowledge that these data can be used to investigate a number of additional questions, such as the specific nature of the encoding of visual orientation in the BOLD signal pattern. It can also be a valuable resource in further optimization of the decoding procedure (classification algorithm, hyper-parameter optimization, etc.). In order to facilitate the required future analyses we have publicly released the data. It has been uploaded to OpenFMRI (accession number: ds000113c) and is also available without restrictions from `GitHub` https://github.com/psychoinformatics-de/studyforrest-data-multires7t and a description is available in DATA IN BRIEF CITATION. We are hoping that this dataset and manuscript serve as starting point to a series of additional analysis that explore aspects beyond acquisition resolution.

## Materials and methods

### Participants

Seven healthy right-handed volunteers (age 21-38 years, 5 males) with normal or corrected to normal vision were paid for their participation. Before every scanning session, they were provided with instructions for the experiment and signed an informed consent form. The study was approved by the Ethics Committee of the Otto-von-Guericke University.

### Stimuli

Following Swisher et al. (2010), a stimulus comprised two semi-annular patches of flickering sine-wave gratings left and right of a central fixation point on a medium gray background (0.8° −7.6° eccentricity, 160° width on each side with a 20° gap along the vertical meridian, above and below the fixation point, to aid separation of gratings between hemifields). Gratings on each side of the stimulus were independently oriented at either 0°, 45°, 90°, or 135°, with a constant spatial frequency of 1.4 cycles per degree of visual angle corresponding to the center of the screen, a contrast of 100%, and a flickering frequency of 4 Hz with 50% duty cycle. The phase of the gratings was changed at a frequency of 4 Hz and was chosen randomly from 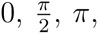 or 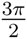 degrees of phase angle (Figure 1).

**Figure 1:**
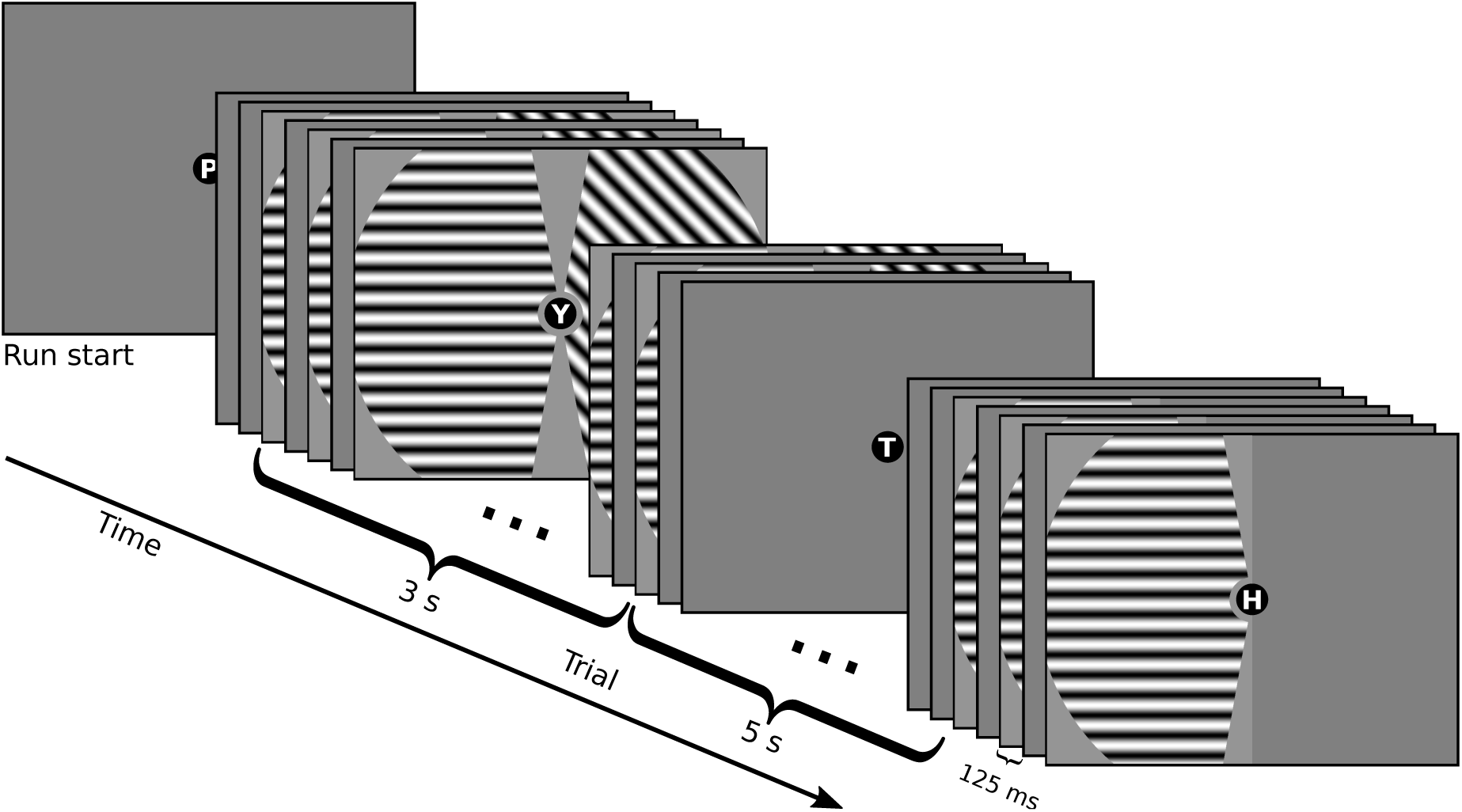
Stimulation paradigm. Independently oriented flickering grating stimuli on a medium gray background were presented in both hemifields for 3 s at the beginning of each trial. Stimulation was followed by a 5 s inter-trial interval. Throughout an entire experiment run, participants performed a continuous central letter reading task to maintain fixation. Interspersed trials where the previous stimulus was repeated in only one of the hemifields were used to decouple stimulation sequences.

Stimulus presentation and response logging were implemented using PsychoPy (v1.79; Peirce, 2008) running on a computer with the (Neuro)Debian operating system (Halchenko and Hanke, 2012). Stimuli were displayed on a rear-projection screen (1280 × 1024 pixels resolution; 60 Hz video refresh rate; 25.5 cm wide) located behind the head coil. Participants viewed the screen via a mirror placed at a distance of ≈4 cm from their eyes. The total viewing distance was 100 cm.

### Behavioral task

In order to keep the participants’ attention focused and to minimize eye-movements, they performed a reading task that was unrelated to the stimulation with oriented gratings. A black circle (radius 0.25°) was presented as a fixation point at the center of the screen. Within this circle, a randomly selected excerpt of song lyrics was shown as a stream of single letters (0.3° height, letter frequency 2 Hz) throughout the entire length of a run. Each trial started with 3 s of stimulation with oriented gratings and continued for another 5 s of a task-only period (Figure 1). During task-only periods, a medium gray background was displayed in both hemifields. At the end of each run, the participant was asked a question related to the previously read text.

In a pilot experiment with in-scanner eye-movement recordings, the letter reading task was found to minimize eye-movements effectively; however, it was unsuitable to verify fixation accuracy on a trial-by-trial basis. In order to evaluate a potential impact of the reading task on the orientation decoding performance, the task was replaced for one participant with a visual detection task. One participant was repeatedly presented with a Landolt C stimulus (radius 0.12°, left or right opening (0.048°) at random intervals in each run. The participants had to respond to the direction of the opening of the probe by pressing one of two buttons corresponding to a left or right opening. Discrimination accuracy for this participant was 98.6%, while orientation decoding performance did not qualitatively differ from mean decoding accuracy of other participants. The performance of the subject with the Landolt C task was compared relative to the 95% binomial proportion confidence interval computed from the number of correct predictions (BOLD pattern classification), concatenated across hemispheres and cross-validation fold, and all subjects performing the reading task. For all resolutions (except 3 mm data) the performance of the subject performing the Landolt C task was within the confidence interval (for 3 mm the decoding accuracy was close to, but higher, than the upper boundary of the confidence interval). This suggests that the employed reading task was generally effective in keeping participants focused on the fixation point.

### Procedures

Participants were scanned in five different sessions, one experiment session for each of the four acquisition resolutions (0.8 mm, 1.4 mm, 2.0 mm and 3.0 mm isotropic) and one session for retinotopic mapping. These sessions took place on different days over the course of five weeks. The order of acquisition resolutions was randomized for each participant. In every experiment session, participants completed ten runs with short breaks in-between, without leaving the scanner. Each run comprised 30 trials (8 s duration; 4 min total run duration). In 20 of these trials, a combination of oriented gratings, one in each hemifield, was presented simultaneously so that each of the four orientations occurred exactly five times in each hemifield. The sequence of orientations was independently randomized for each hemifield, resulting in random pairings of orientations within trials. In order to decouple stimulation sequences between hemifields, ten NULL events were inserted into the trial sequence at pseudo-random positions (a run could not start with a NULL event and no two NULL events could occur in immediate succession). NULL events were identical to regular trials, except for the fact that in one hemifield the same oriented grating as in the previous trial was repeated while the other hemifield remained empty. The side of the blank hemifield was chosen at random for each NULL event. For all participants, the actual generated trial sequences show a roughly equal count of NULL events for each hemifield and frequency of unique combinations of grating orientations (refer to supplementary material section *Experimental Design* for more details).

### Functional imaging

The objective for functional data acquisition was to obtain BOLD fMRI data from the V1 ROI at four different resolutions with an identical stimulation paradigm. MR acquisition parameters were chosen to be maximally similar across resolutions given two a priori constraints: 1) sufficient spatial coverage of the V1 ROI and 2) identical sampling frequency (TR) across resolutions.

T2*-weighted echo planar images (EPI) (TR/TE=2000/22 ms, FA=90°) of the occipital lobe were acquired during visual stimulation using a 7 Tesla whole body scanner (Siemens, Erlangen, Germany) and a 32 receive channel head coil (Nova Medical, Wilmington, MA). Slices, oriented parallel to the calcarine sulcus (on a tilted axial plane), were acquired for 4 different spatial resolutions: 3 mm isotropic (FoV=198 mm, matrix size 66 × 66, 37 slices, GRAPPA accel. factor 2), 2 mm isotropic (FoV=200 mm, matrix size 100 × 100, 37 slices, GRAPPA accel. factor 3), 1.4 mm isotropic (FoV=196 mm, matrix size 140 × 140, 32 slices, GRAPPA accel. factor 3) and 0.8 mm isotropic (FoV=128 × 166.4 mm (AP × LR), matrix size 160 × 208, 32 slices, GRAPPA accel. factor 4). All EPI scans implemented ascending slice acquisition order and used a 10% inter-slice gap to minimize cross-slice excitation. For example, for a 3 mm acquisition, the acquired voxel dimension was 3× 3 × 3 mm, plus a 0.3 mm interslice gap. The sequence for 0.8 mm resolution used a left-right phase encoding direction in order to avoid wrap-around artifacts, while all other sequences used anterior-posterior phase encoding. 121 volumes were acquired for each experiment run and 10 separate scans (one for each experimental run) were performed for each subject. An automatic positioning system (Siemens AutoAlign Head LS) was used to aid positioning of the field-of-view to the same volume in each scan for each subject similar to the procedure described in Dou et al. (2014). Online distortion correction (In and Speck, 2012) was applied to data from all the scans.

As a result of the technical constraints the scan volume of the 0.8 mm acquisitions was substantially smaller than that of the other resolutions and did not cover all of the V1 ROI. In order to aid co-registration of the small scan volume with the structural image, an additional EPI acquisition was performed that used the same auto-alignment procedure, but with a 250 × 250 in-plane matrix and 57 slices (4 s TR). This setup increased the FoV in the axial plane to cover the full extent of the brain, while the 20 additional slices further increased the coverage along the inferior-superior direction. 60 volumes were acquired to improve image signal-to-noise ratio (SNR) by averaging across volumes. The resulting volume was used as an intermediate alignment target.

Figure S7 illustrates the effect of distortion correction and the alignment quality of BOLD images to the respective structural images for two participants.

### Structural imaging

Structural images and susceptibility weighted images were acquired for all participants in a 3 Tesla Philips Achieva equipped with a 32 channel head coil using standard clinical acquisition protocols. T1-weighted image consisted of 274 sagittal slices (FoV 191.8 × 256 × 256 mm) and an acquisition voxel size of 0.7 mm with a 384 × 384 in-plane reconstruction matrix (0.67 mm isotropic resolution). It was recorded using a 3D turbo field echo (TFE) sequence (TR 2500 ms, inversion time (TI) 900 ms, flip angle 8°, echo time (TE) 5.7 ms, bandwidth 144.4 Hz/px, SENSE reduction AP 1.2, RL 2.0). A 3D turbo spin-echo (TSE) sequence (TR 2500 ms, TE eff 230 ms, strong SPIR fat suppression, TSE factor 105, bandwidth 744.8 Hz/px, SENSE reduction AP 2.0, RL 2.0, scan duration 7:40 min) was used to acquire a T2-weighted image whose geometric properties otherwise match the T1-weighted image. A susceptibility weighted image with 500 axial slices (thickness 0.35 mm, FoV 181× 202× 175 mm) and an in-plane acquisition voxel size of 0.7 mm reconstructed at 0.43 mm (512× 512 matrix) was recorded using a 3D Presto fast field echo (FFE) sequence (TR 19 ms, TE shifted 26 ms, flip angle 10°, bandwidth 217.2 Hz/px, NSA 2, SENSE reduction AP 2.5, FH 2.0). All the acquisition protocols used for recording anatomical images and susceptibility images were identical to those used in Hanke et al. (2014).

### Region of interest localization

Standard retinotopic measurements were performed using four runs of stimulation with flickering checkerboard patterns (Warnking et al., 2002), one run each for contracting/expanding rings and clockwise/counter-clockwise wedges. During stimulation, participants fixated the center of the screen while performing the letter reading task described above. Each run comprised five stimulus cycles, plus 4 s and 12 s of task-only periods (no checkerboard stimulus) at the start and at the end of a run respectively. fMRI acquisition took place in the same 3 Tesla scanner as the structural imaging. Full brain acquisition was performed with T2*-weighted gradient echo, single-shot echo planar imaging (EPI) sequence (TR/TE=2000/30 ms, FA=90°, SENSE reduction AP 2) with 3 mm isotropic voxel size, and 10% inter-slice gap (FoV=240 mm, matrix size 80× 80, 35 slices, ascending order, anterior-to-posterior phase encoding direction). 90 volumes were acquired in each run.

Retinotopic phase maps (polar angle and eccentricity) were generated using the 3DRetinophase tool in the AFNI software package (Cox, 1996). The V1 region was manually delineated on the cortical surface (following the procedure described in Warnking et al., 2002). Surface reconstruction was performed using the default Freesurfer recon-all pipeline (Dale et al., 1999), using T1 and T2-weighted images as input. V1 delineations on the surface were projected back into a subject’s individual volumetric space to generate a participant specific V1 ROI mask for the classification analyses. Figure S7 demonstrates the alignment of the 7T BOLD fMRI with the reconstructed cortical surface.

The associated raw data are available is part of dataset ds000113d on OpenFMRI and are further described in Sengupta et al. (2016).

### Blood vessel localization

A volumetric mask of V1 voxels with venous contributions was generated for each subject using the following procedure. First, the phase component of the SWI scan was masked (using a brain masked derived from the magnitude component), and 3D phase unwrapped with PRELUDE (default settings; Jenkinson, 2003) from FSL (v5.0.8; Smith et al., 2004). Following the procedure outlined in Haacke et al. (2004), the unwrapped phase image was spatially high-pass filtered using a mean ‘box’ filter kernel (65x65x65 voxels, as implemented in fslmaths; Smith et al., 2004). The high pass filtered phase component *ϕ*(*x*) was then transformed to a score *g*(*x*) (value interval [0, 1]) using *g*(*x*) = (*π – φ*(*x*))*/π* for 0 *< φ*(*x*) ≤ *π* and 1 otherwise. These scores were multiplied 4 times with the original magnitude image, as suggested by Haacke et al. (2004), in order to enhance the contrast between venous and non-venous voxels. Blood vessel masks computed from the thresholded enhanced magnitude image were resliced into different acquisition resolutions using trilinear interpolation and were constrained to individual V1 masks for each participant.

Separate MVP analyses were performed inside and outside the venous voxels (with variable mask intensity threshold) in V1 to investigate their individual contributions at different acquisition resolutions across different threshold levels.

The associated raw data are available is part of dataset ds000113 on OpenFMRI and are further described in Hanke et al. (2014).

### Orientation decoding analysis

MVP analysis for orientation decoding was performed with PyMVPA (v2.4.1; Hanke et al., 2009) on a compute cluster running (Neuro)Debian (v8.0; Halchenko and Hanke, 2012). For feature extraction, BOLD fMRI time series from an individual experimental run were voxel-wise fitted to hemodynamic response (HR) regressors (boxcar function convolved with the canonical Glover HRF kernel (Glover, 1999) for each experimental condition using a general linear model (GLM). Additionally, the GLM design matrix included temporal derivatives of HR regressors, six nuisance regressors for motion (translation and rotation), and polynomial regressors (up to 2nd-order) modeling temporal signal drift as regressors of no-interest. GLM *β* weights were computed using the GLM implementation in NiPy (v0.3; Millman and Brett, 2007) while accounting for serial correlation with an autoregressive term (AR1). Lastly, separately for every run *β* scores were *Z*-scored per voxel. The resulting dataset for MVP analysis contained 40 samples (one normalized *β* score per condition per run) for each participant.

Linear support vector machines (SVM; PyMVPA’s LinearCSVMC implementation of the LIBSVM classification algorithm; Chang and Lin, 2011) were used to perform a within-subject leave-one-run-out cross-validation of 4-way multi-class orientation classification. This method was selected based on its prevalence in the literature, not because of an assumed optimal performance in this context. This linear SVM algorithm has one critical hyper-parameter *C* that indicates the trade-off between width of the margin of the classifying hyperplane and number of correctly classified training data points. While it seems uncommon for neuroimaging studies to optimize this parameter for a particular application, we observed substantial variability in performance with varying number of input features. Consequently, we decided to tune this parameter using a nested cross-validation approach, where the training portion within each cross-validation fold was subjected to a series of leave-another-run-out cross-validation analyses in order to perform a grid search for the optimal *C* value (search interval [10^−5^, 5 ×10^−2^] in 200 equal steps). The “optimal” *C* value was then used to train a classifier on the full training dataset, which was subsequently evaluated on the data from the left out run. Reported accuracies always refer to the performance on the test dataset using the tuned *C* setting. Tuning of the *C* parameter was performed independently for each participant, resolution, and hemisphere. The ranges of tuned *C* parameters for all resolutions are illustrated in Figure S6.

### Spatial filtering strategies

In order to investigate how signal for orientation decoding is distributed across the spatial frequency spectrum, two different strategies for volumetric spatial filtering of the functional imaging data were implemented.

#### Gaussian smoothing

Similar to Swisher et al. (2010), we used Gaussian filtering prior feature extraction for MVP analysis to estimate the spatial scale of the orientation specific signal. In the following, the size of the Gaussian filter kernel is described by its full width at half maximum (FWHM) in mm. Individual filters were implemented using the following procedure: *Low-pass* (LP) 3D Gaussian spatial filtering was performed with the image smooth() function in the nilearn package (Pedregosa et al., 2011). *High-pass* (HP) filtered images for a particular filter size were computed by subtracting the respective LP filtered image from the original, unfiltered image. *Bandpass* (BP) filtering was implemented by a Difference-of-Gaussians (DoG) filter (Alink et al., 2013). Filtered images were computed by subtracting the LP filtered images for two filter sizes from each other. For example, an image for the “4-5 mm” band was computed by subtracting the 5 mm LP filtered image from the 4 mm LP filtered image. It is important to note that, due to the nature of the filter, the pass-band of a DoG filter is not as narrow as the filter-size label might suggest. Figure S5 illustrates the attenuation profile of an exemplary 4-5 mm DoG filter. However, for compactness and compatibility with previous studies (*e.g.*, Alink et al., 2013) we are characterizing DoG BP filters by the FWHM size of the underlying LP filters. The respective *band-stop* (BS) filtered image were computed by subtracting the corresponding BP filtered image from the original, unfiltered image.

Because of its prevalence in standard fMRI analysis pipelines, spatial filtering was always applied to the whole volume, prior to any masking. However, as this procedure leads to leakage of information from outside the ROI into the ROI due to smoothing, particularly with large-sized LP filters, we also performed a supplementary analysis where filtering was restricted to the V1 ROIs in each hemisphere to prevent information propagation by smoothing (see supplementary material).

#### Spatial resampling to other resolutions, with and without Gaussian filtering

A frequently expressed concern in the literature with respect to Gaussian smoothing is that a linear transformation does not actually remove high spatial frequency information (Alink et al., 2013; Kamitani and Sawahata, 2010); instead, it merely implements a relative scaling of frequency components (see Misaki et al., 2013). In order to explore any potential impact of an irreversible frequency-domain transformation, we performed a Fourier (FFT) based spatial frequency resampling, which destructively removes high-frequency components, using the scipy function signal.resample() (Jones et al., 2001). For details on the procedure see the supplementary material. The V1 ROI masks were linearly interpolated into the resampled space with the ndimage.interpolation.zoom() function in scipy. FFT resampling was also combined with subsequent Gaussian lowpass filtering in order to evaluate a suggestion by Freeman et al. (2013) that one way of testing the contribution of fine scale signals to orientation decoding is to compare high-resolution BOLD fMRI data down-sampled to conventional resolutions, with or without first removing high spatial frequency signals. For all spatial resampling analysis, with or without Gaussian filtering, all voxels in the respective V1 ROI masks were considered for multivariate decoding.

## Results

### Decoding performance on native acquisition resolution

#### Effect of acquisition resolution and number of input voxels

In order to determine the effect of acquisition resolution, we performed orientation decoding at all resolutions. Figure 2A shows the mean classification accuracy across participants and hemispheres as a function of acquisition resolution in the V1 ROI. In the set of tested acquisition resolutions, we found the peak classification performance of 40.89% at 2 mm isotropic resolution.

**Figure 2:**
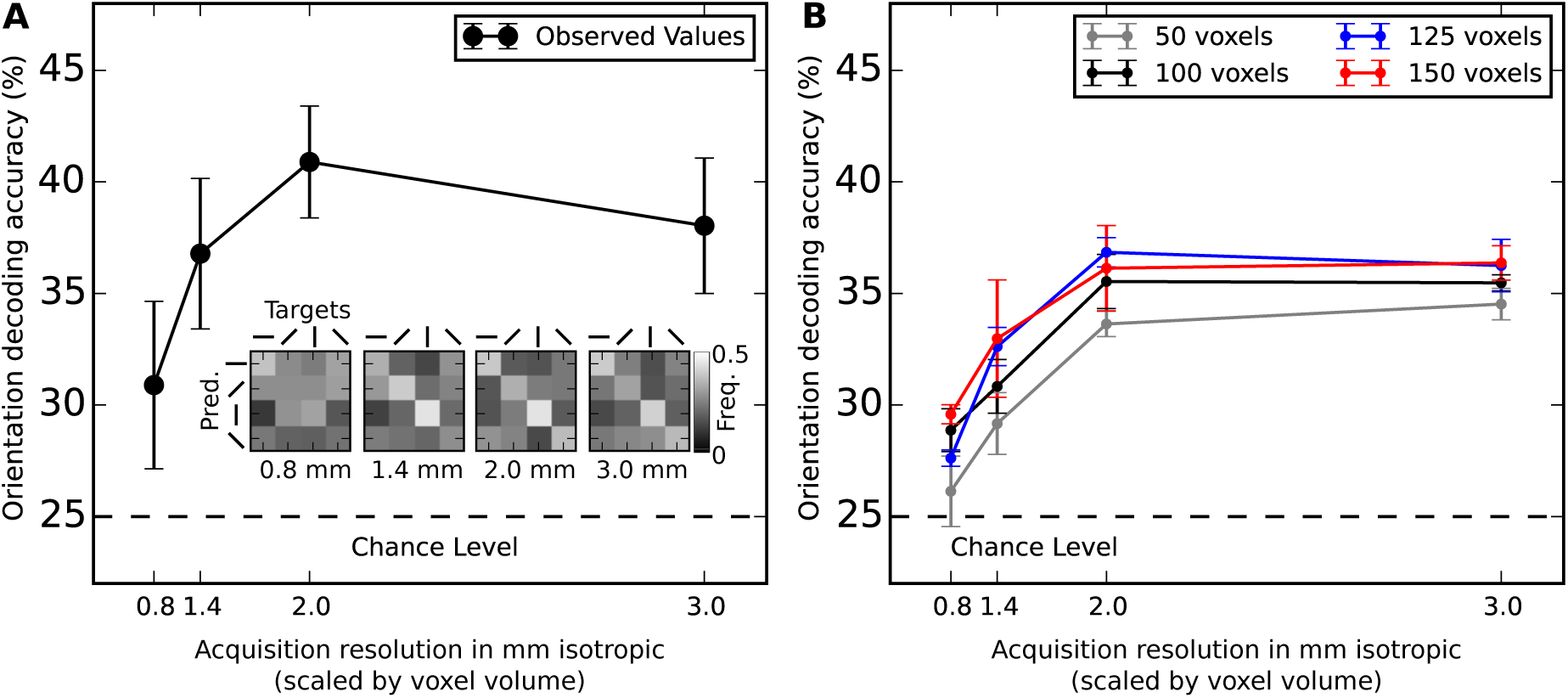
**(A)** Orientation decoding accuracy on spatially unfiltered data as a function of acquisition resolution in the whole contralateral V1 ROI. Error bars show the standard error of the mean (SEM) across 7 participants averaged across hemispheres. Chance level accuracy (25%) is indicated as a horizontal dashed line. Classification performance is detailed in confusion matrices for each resolution depicting the frequency of correct classification for each combination of prediction and target values. **(B)** Analog to (A), but with a constant number of input voxels across resolutions. 50, 100, 125, or 150 voxels were selected at random from the the whole contralateral V1 ROI for the classification analysis. Selection was repeated 100 times. Error bars show SEM across repetitions. Upper range limit of 150 voxels was determined by the ROI with the least number of voxels at 3 mm resolution.

For the above analysis, all voxels in the respective V1 ROIs were used. As the number of voxels in a 0.8 mm V1 mask was substantially higher than those in a 3.0 mm V1 mask (Table 1) and the number of input features/voxels can impact the classification performance, we repeated the analysis, but held the number of voxels constant across participants and resolutions (50, 100, 125, and 150 voxels). Voxel sub-selection was done randomly, and the analysis was repeated 100 times with a new random selection of voxels. Figure 2B shows that a constant and smaller number of input voxels had a negative effect on classification performance. Classification performance was better with 2.0 mm and 3.0 mm data as compared to 0.8 mm and 1.4 mm data.

**Table 1:** V1 ROI size. Average number of voxels for both hemispheres with standard deviation across participants. The four rightmost columns indicate the number of voxels within the ROI that are considered to be intersecting veins for two different thresholds (the 40% of voxels with the highest volume fraction of blood vessels; and the same for the top 10% voxels; see Figure 6 for an illustration).

| Resolution | V1 Region of Interest |  |  |  | Venous voxels in V1 for two thresholds |  |  |  |
| --- | --- | --- | --- | --- | --- | --- | --- | --- |
|  | Left hemisphere |  | Right hemisphere |  | >60<sup>th</sup> percentile |  | >90<sup>th</sup> percentile |  |
|  | #voxels | std | #voxels | std | #voxels | std | #voxels | std |
| 0.8 mm | 7312 | 1912 | 7683 | 2556 | 1148 | 446 | 287 | 111 |
| 1.4 mm | 2084 | 626 | 2169 | 710 | 518 | 186 | 130 | 47 |
| 2.0 mm | 883 | 273 | 898 | 311 | 231 | 84 | 58 | 21 |
| 3.0 mm | 324 | 94 | 327 | 104 | 105 | 36 | 26 | 9 |

**Figure 6:**
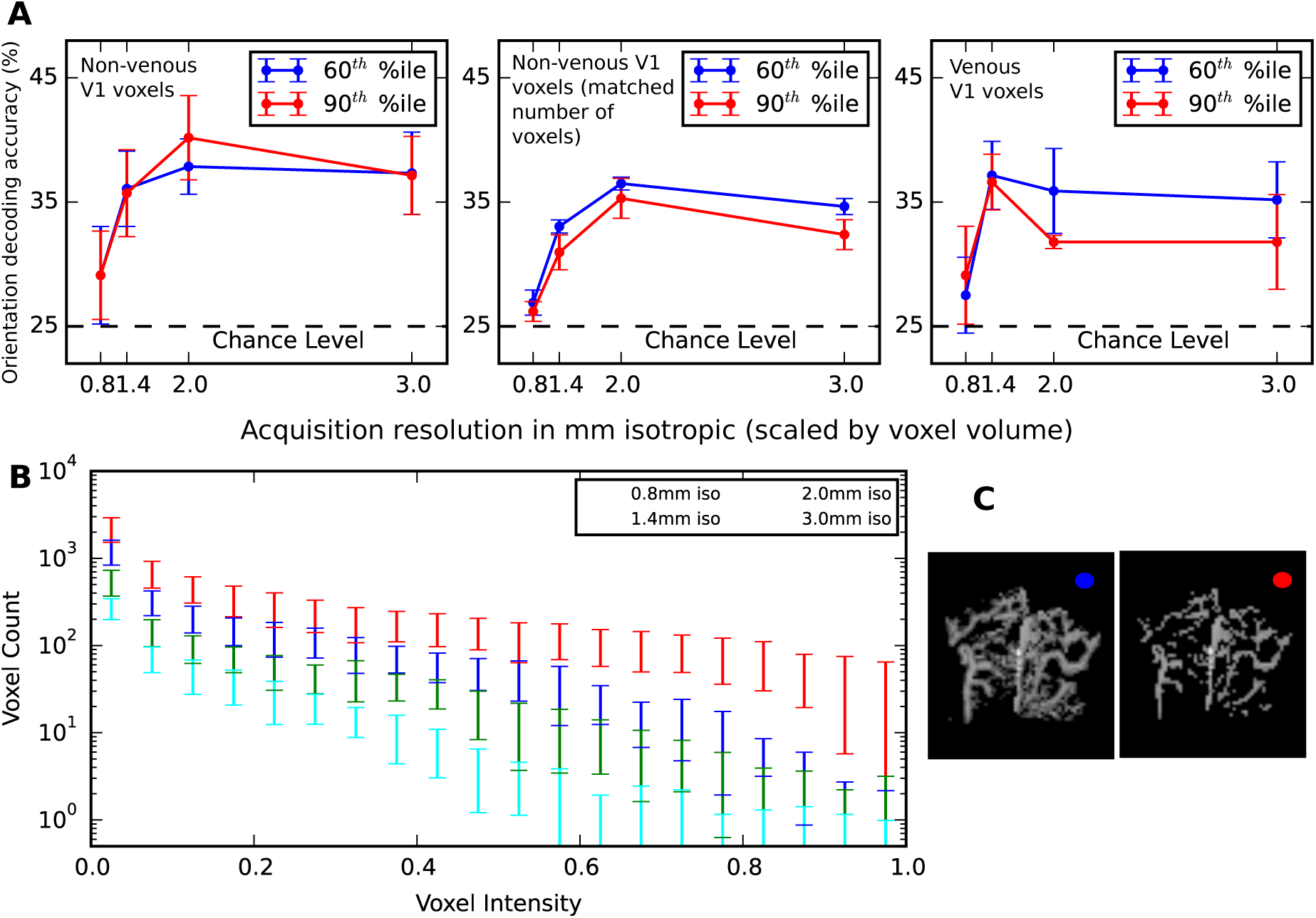
**(A)** Decoding accuracy computed inside and outside the vein mask within the V1 ROI. The vein masks obtained from susceptibility weighted imaging were thresholded at two different levels i.e. 60 percentile and 90 percentile. The panel on the left shows the performance of the entire V1 ROI outside the vein mask (non-venous voxels) for the two different thresholds. Orientation decoding accuracy on V1 voxels restricted to the veins mask (venous voxels) is shown on the right panel. The middle panel depicts the average decoding performance of non-venous voxels that randomly sampled and matched in number to the corresponding venous voxels. The dashed horizontal lines indicate the chance performance. **(B)** Trilinear interpolation was used to reslice the vein mask to all four target resolutions. The histogram shows the distribution of mask voxel intensities corresponding to the volumetric fraction of “vein voxels” in the high-resolution vein mask (voxel count axis in log-scale). **(C)** Axial maximum intensity projection of the vein mask of one participant resliced to the 0.8 mm resolution; illustrates the two chosen thresholds. The color indicator correspond to the curves depicted in panel A.

#### Time-series signal-to-noise ratio (tSNR)

It has been shown that overall contrast-to-noise ratio (OCNR) is a factor that impacts classification performance (Chaimow et al., 2011). According to Chaimow et al. (2011) OCNR is proportional to contrast range and the square root of the number of voxels and is inversely proportional to the noise level. The noise level was calculated as the inverse of time course signal-to-noise ratio, which in turn depends on voxel size (Triantafyllou et al., 2005). In this study, tSNR is modulated across acquisition resolutions due to differential impact of technical/thermal and physiological noise components. In order to characterize this impact, we computed tSNR for each voxel as the ratio of mean signal intensity across all time points after polynomial detrending (1^st^ and 2^nd^ order; analog to preprocessing for MVP analysis) of scanner drift noise and the corresponding standard deviation. Voxel-wise tSNR was averaged across all experiment runs. For a tSNR estimate of the whole ROI, we averaged this score across all voxels. The relationship of voxel volume and tSNR in the empirical data can be well explained by the following model (Triantafyllou et al., 2005):

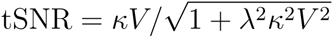

where *V* is the voxel volume, *κ* is the proportionality constant, and *λ* is the magnetic field strength independent constant parameter with *λ*=0.0117, *κ*=22.74 (*R*^2^=0.95) The estimated asymptotic tSNR limit of 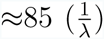 is similar to the report of Triantafyllou et al. (2005) for 7 Tesla acquisitions and is reached around 2.5 mm acquisition resolution (see supplementary Figure S8).

Figure 3A illustrates the non-linear relation of tSNR and orientation decoding accuracy. We observe a substantial drop in accuracy when decreasing resolution from 2 mm to 3 mm, despite a further increase in tSNR. This non-linearity was not observed by Gardumi et al. (2016), who only reported a positive trend for the correlation between decoding accuracy and tSNR, based on a single acquisition (1.1 mm resolution with comparable tSNR of ≈32, and other resolutions being generated by reconstructing k-space data to lower resolutions).

**Figure 3:**
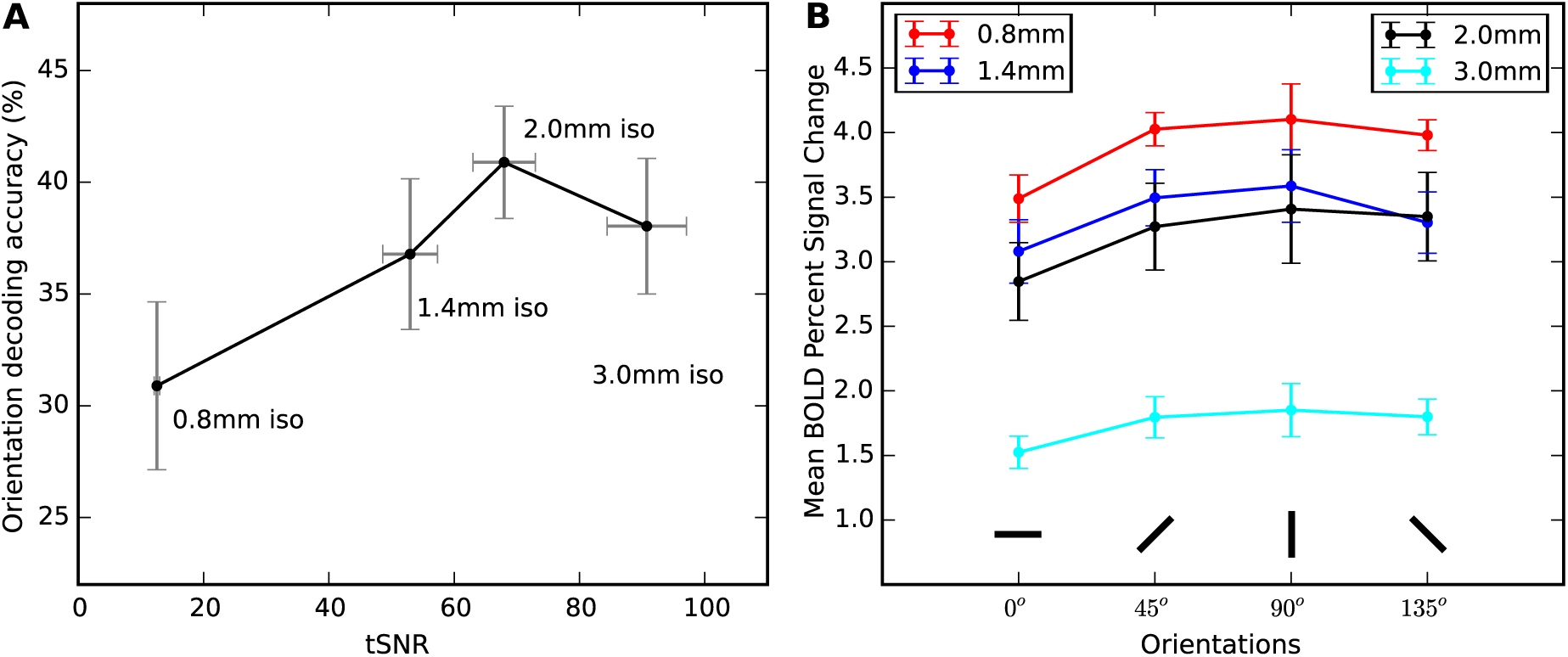
**(A)** Orientation decoding accuracy vs. temporal signal-to-noise ratio (tSNR) for the four measurements (resolutions). Error bars show the SEM for tSNR and accuracy across subjects and hemispheres. **(B)** Estimated BOLD signal change by orientation for all resolutions. Maximum pairwise signal change difference is observed for the cardinal directions 0° and 90°. This pattern is congruent with the confusion plots in Figure 2A.

#### BOLD signal change

Another potential source of differences in orientation decoding accuracy across resolutions are BOLD signal amplitude differences due to, for example, differential impact of a partial voluming effect (see Tong et al., 2012; Alink et al., 2013). In order to quantify this effect, we calculated mean percentage BOLD signal change in response to any flickering orientation stimulus across resolutions using FeatQuery in FSL (v5.0.8; Smith et al., 2004). Similar to preprocessing in MVP analysis, no spatial smoothing was performed before calculating the percentage signal change. In order to obtain comparable percentage signal change across resolutions, we obtained a mask of all responsive V1 voxels (*z >* 2.3 with *p <* 0.05 default parameters of FSL FEAT) in 0.8 mm data for every subjects (Swisher et al., 2010, similar to Figure 3). The responsive V1 voxel mask obtained at 0.8 mm was resliced into 1.4 mm, 2.0 mm and 3.0 mm resolutions. Percentage signal change was calculated with FeatQuery within these masks. We found that the mean percentage BOLD signal change was the highest for 0.8 mm resolution: 4.51% (0.8 mm), 3.92% (1.4 mm), 3.73% (2.0 mm), and 2.05% (3.0 mm).

Previous studies found differential BOLD response magnitudes to different visual orientations. Furmanski and Engel (2000) reported stronger responses to cardinal orientations. In contrast, Swisher et al. (2010) found greater responses to oblique orientations. In order to test for a differential effect and a possible interaction between orientation and acquisition resolution, we computed a 2-factor (orientation and resolution) within-subject ANOVA for the estimated BOLD signal change from all 7 subjects (Figure 3B). There was a significant main effect of acquisition resolution (*F* (3, 18) = 32.99, *p <* 0.001). We found no statistically significant main effect of orientation (*F* (1.22, 7.319) = 4.678, *p* = 0.061; using Greenhouse-Geisser correction due to violation of sphericity assumption, Mauchly’s test *p* = 0.002). There was a non significant interaction between the factors, resolution, and orientation (*F* (2.947, 17.68)=1.96, p=0.158 after Greenhouse-Geisser correction).

#### Impact of head motion on decoding accuracy

Head motion is a likely factor to impact decoding accuracy. In order to evaluate this effect, we calculated a head motion index suggested by Alink et al. (2013) for every participant and acquisition resolution. Inline with the findings of Gardumi et al. (2016), we found a consistent, but non-significant trend towards a negative correlation between head motion and decoding accuracy across acquisition resolutions 0.8 mm: r=-0.45, p=0.30; 1.4 mm:r=-0.64, p=0.11; 2.0 mm: r=- 0.68, p=0.09; 3.0 mm: r=-0.23, p=0.60).

### Decoding performance on spatially filtered data

#### Impact of Gaussian smoothing

Figure 4 A-D show the impact of Gaussian filtering on the classification performance for data from all four acquisition resolutions. LP spatial filtering is most commonly performed as a noise reduction step in fMRI data pre-processing. The classification performance achieved on HP filtered data of the same filter size is an indication of the amount of usable information removed by LP filtering. Classification performance on BP filtered data indicates whether usable information is present in a particular band of spatial frequencies. Likewise, band-stop performance indicates the presence of usable information anywhere, except in a particular band.

**Figure 4:**
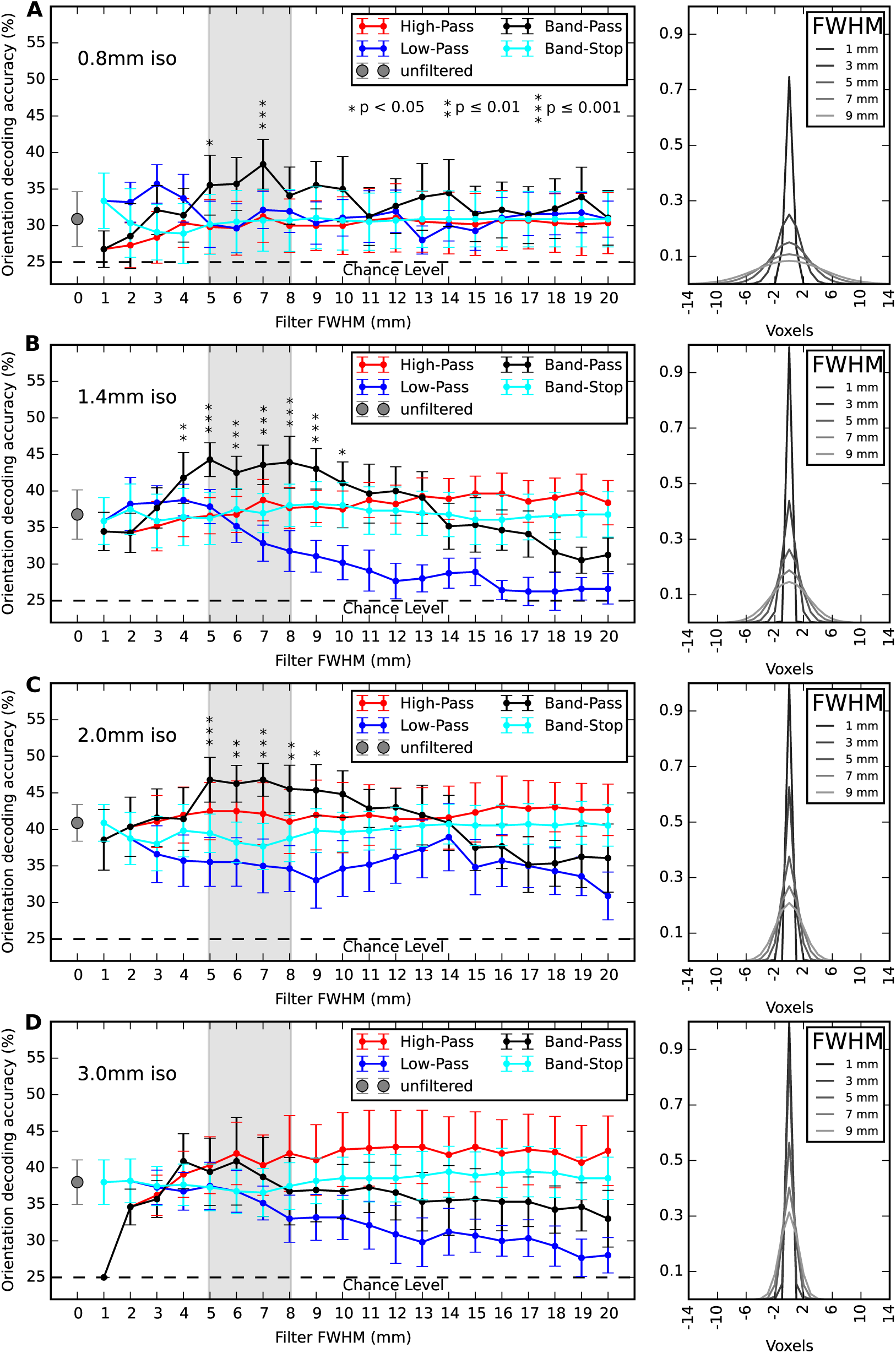
Orientation decoding accuracies for all acquisition resolutions (increasing acquisition voxel size from top to bottom) and levels of spatial high-pass, low-pass, band-pass, and band-stop Gaussian filtering. Panels on the right visualize the size of selected Gaussian filter kernels with respect to the voxel size at each resolution. FWHM values for band-pass and band-stop filters refer to the corresponding 1 mm band to the closest smaller filter size (*e.g.*, 5 mm refers to the 4-5 mm band). McNemar test (Edwards, 1948) was used to comparing the performance of the BP filtered data with the unfiltered decoding performance. Star markers indicate a significant difference (Bonferroni-corrected, see legend in (A) for criteria).

Except for 0.8 mm and 1.4 mm data, we observed no increase in mean decoding for LP filtering, compared to performance on unfiltered data. For all resolutions, except for 0.8 mm, we see observe the best performance after LP filtering with kernel sizes no larger than 3 mm FWHM. Peak performance on HP filtered data was achieved for filter sizes larger than 9 mm FWHM, except for the 0.8 mm acquisition resolution. We investigated via BP filtering which frequency bands were most informative for orientation decoding across all acquisition resolutions, using DoG BP filters with a 1 mm difference in the FWHM size of the underlying Gaussian filters (Figure 4 A-D; black curves). The results show peak performance of BP filtering yielded for all acquisition resolutions in the range of 5-8 mm (highlighted range).

Average decoding accuracy of BS filtered data remained above-chance for all spatial frequency bands. The BS performance curve initially follows the LP performance for small filter sizes, but resembles the HP performance for larger filter sizes.

#### Impact of spatial resampling to other resolutions, with and without Gaussian smoothing

As an alternative approach to Gaussian LP filtering for simulating a resolution reduction, data acquired in a particular resolution were resampled (FFT-based transformation) to all other resolutions and classification analysis was performed with and without additional prior Gaussian LP filtering, as suggested by Freeman et al. (2013). The results are depicted in Figure 5.

**Figure 5:**
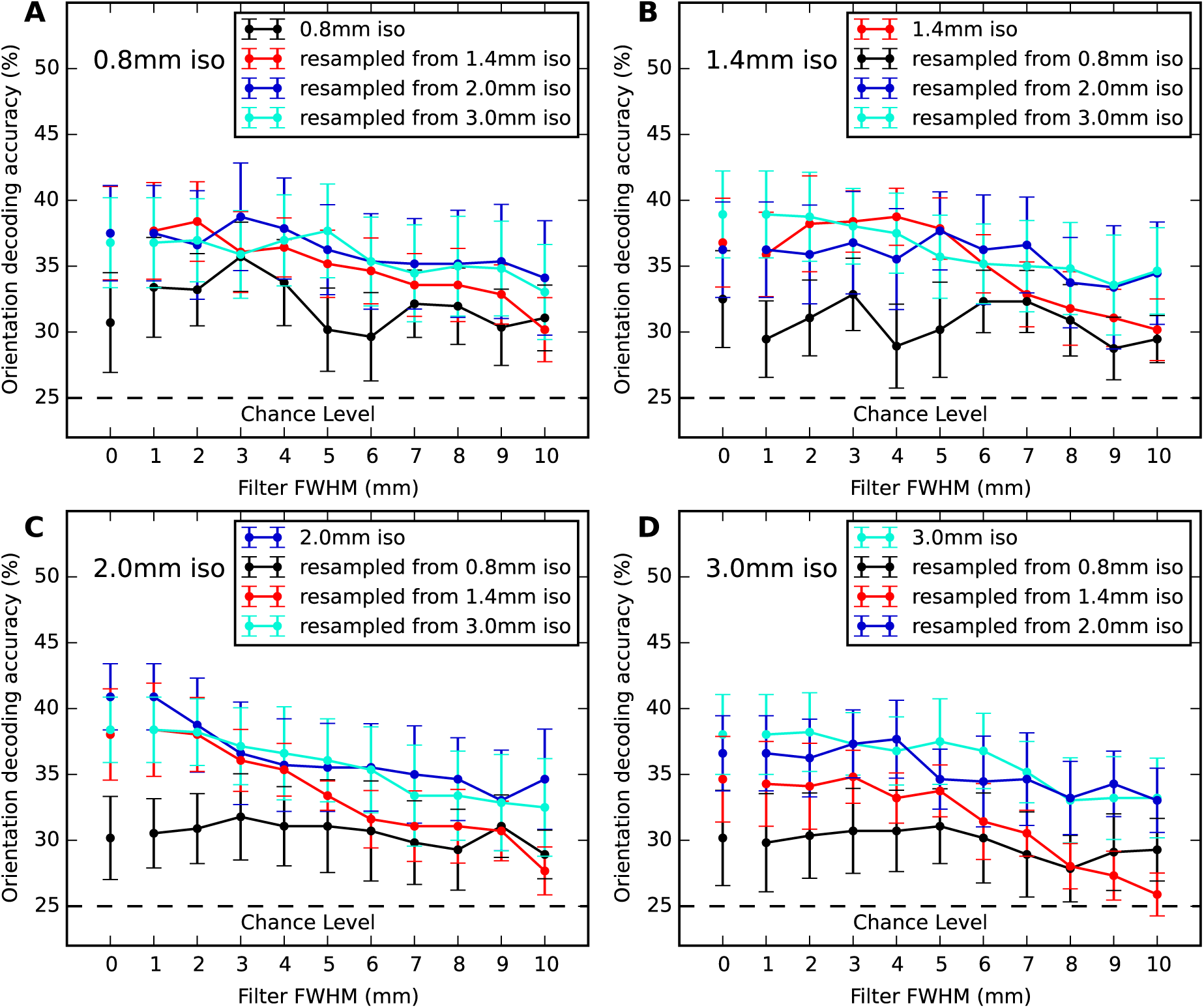
Orientation decoding performance on fMRI data resampled to other spatial resolutions, with and without different levels of prior low-pass Gaussian spatial filtering. Each panel title indicates the respective resolution after spatial resampling. The panel legends identify the corresponding original native acquisition resolutions. The color coding consistently identifies the native resolution across all four panels. The disconnected data points at 0 mm represent the decoding accuracy after spatial resample *without* prior Gaussian LP filtering. Recording high-resolution data with subsequent spatial down-sampling tends to yield lower classification accuracies compared to the native resolution acquisition, with or without prior Gaussian low-pass filtering of any tested kernel size.

We observed no general benefit of spatial down-sampling with respect to decoding accuracy. We also did not see systematically improved accuracies after LP filtering across resolutions.

Data acquired at 2.0 mm or 3.0 mm showed a general trend towards higher decoding accuracy after resampling (up-sampling or down-sampling) compared to the corresponding native acquisition resolution, even with prior Gaussian LP filtering of different kernel sizes. 0.8 mm data consistently showed low decoding accuracy when resampled to any other resolution with or without Gaussian filtering.

### Vascular contribution to orientation decoding

Orientation decoding again performed inside and outside the vein localizer mask in the V1 ROI in order to evaluate the availability of orientation discriminating signal in the vascular system. Two different, arbitrary thresholds were used to classify voxels as intersecting vs. non-intersecting with veins, based on the co-registered and re-sliced vein mask: the top 40% and top 10% of voxels with the highest value after realignment and reslicing to the target resolution with trilinear interpolation. The resulting number of voxels are presented in Table 1.

Analyses outside the vein mask were performed twice: once for the entire region and again for a subset of voxels that was constrained to the number of voxels inside the vein mask for the corresponding resolution. In the latter case, the analysis was repeated with a new random voxel selection 100 times.

Figure 6A (right panel) shows that voxels with the highest venous content in their volume still yield above chance decoding performance. The performance drop for the two lowest resolutions between the two vein mask thresholds may be explained by the low number of input features going into the classification at high threshold (compare Figure 6A, left panel). At 0.8 mm, the 10% most venous voxels yield the same decoding performance as the rest of the V1 ROI combined (Fig. 6, middle panel), and noticeably more than a corresponding number of randomly samples non-venous voxels (Fig. 6A, middle panel). Similar results can be observed for the 1.4 mm resolution.

## Discussion

In order to explore the effect of acquisition resolution and spatial filtering on the decoding of visual orientations from primary visual cortex, we measured ultra-high field 7 Tesla fMRI data at four different resolutions from seven participants. Linear SVM classifiers were trained to classify voxel patterns of regression weights of hemodynamic response models for the visual stimulation with four different oriented gratings. Cross-validated classification accuracy was used as performance metric.

The overall classification accuracies reported here are deceptively low (peaking at 40-50% with a theoretical chance-level performance of 25% for the 4-way classification analyses employed in this study). Other decoding studies in the literature have often used binary classification paradigms (for example, Alink et al., 2013; Chaimow et al., 2011) or reported average pairwise accuracy for classification performance results like (*e.g.*, Kamitani and Tong, 2005; Op de Beeck, 2010). Converted into average pairwise binary accuracies, the results reported here range from 55% to 70% (for 0.8 mm and 2 mm respectively, each accuracy corresponding to an analysis of the full V1 ROI and with no additional smoothing; see Figure 2A; theoretical chance-performance: 50%), hence accuracies are of the same magnitude as in other studies (see, for example, Haynes and Rees, 2005; Alink et al., 2013). In addition, some studies like Swisher et al. (2010) also reported similar unfiltered accuracy results (≈ 50%) in a 4-way classification analysis with 0°, 45°, 90°, and 135° gratings with much longer stimulation time (a block design of 18s of block duration and 8 blocks/run). Therefore, we conclude that the overall quality of the present data is comparable to that of previous studies, and that the results presented here can be used to address open questions regarding the impact of data acquisition and spatial filtering parameters on the decoding of orientation from the early visual cortex. Importantly, in this experiment we did not use a univariate feature selection approach to define the “visually responsive” voxels in V1 (for example a GLM contrast) in an attempt to improve the decoding accuracy. Studying the potential impact of such an approach is left to a future study.

Given the uncertainty of how much a further optimization of the decoding procedure – *e.g.*, the specific classification algorithm, hyper-parameter optimization strategy, and feature selection method – would impact the results, we consider the interpretation of the present results regarding the nature of the signal source as a starting point for a further exploration of their robustness respect to variations of analysis parameters not considered here.

### Optimal acquisition resolution

Among the four tested acquisition resolutions, the highest decoding accuracy was observed with a 2 mm resolution (Figure 2A). This result is congruent with a simulation study by Chaimow et al. (2011) that analyzed the impact of anatomical and physiological properties of primary visual cortex, as well as technical parameters of BOLD fMRI acquisition on the accuracy of decoding the stimulated hemifield from signal sampled from ocular dominance columns. The aforementioned study included a number of predictions for choosing optimal voxel size and number of input voxels to maximize decoding accuracy for 3 Tesla fMRI (see Figure 6 in Chaimow et al., 2011) that show a striking similarity to the results presented here (Figure 2). For 3 Tesla fMRI, Chaimow et al. (2011) showed that peak decoding accuracy is achieved around 3 mm in-plane voxel size for ocular dominance. Given that the profile of orientation columns has higher spatial frequency compared to ocular dominance columns (Obermayer and Blasdel, 1993) and the BOLD PSF at 7 Tesla is considerably smaller compared to 3 Tesla (Shmuel et al., 2007; Engel et al., 1997) a higher optimal resolution was to be expected for this study, and this hypothesis is supported by our results. This finding is also inline with a recent study by Gardumi et al. (2016) showing that optimal decoding accuracy of speaker identity, or phonemes, from auditory cortex BOLD patterns could be achieved with an effective voxel size of 2.2 mm (acquisition resolution was 1.1 mm and target resolution was achieved by reconstructing k-space data to a lower resolution), although the nature of the commonality between these findings remains to be investigated.

Superior decoding performance at 2 mm was still observed even when the number of input voxels for classification was held constant across resolutions, although the performance differences between resolutions are reduced (Figure 2B). The ratio of input features (voxels) and the number of observations (fixed in this study) is a critical factor for the training of a classification model, as with increasing dimensionality the sampling of the feature space becomes sparser, and, consequently, the estimated decision surface suffers from increased uncertainty (curse of dimensionality, Bellman, 1961, after Friedman et al. 2001) In this study, the number of voxels in the ROI varies by a factor of >20 from the lowest to the highest resolution (Table 1) and the coverage volume also varies across resolutions. Despite full V1 coverage for the 1.4 mm (except for one subject), 2 mm, and 3 mm acquisitions, peak decoding accuracy was observed with 2 mm data. There is no noticeable difference between accuracies for the lowest resolutions when the number of input voxels is held equal (Figure 2B). The pattern of decoding accuracy differences when using the full ROI vs. a constant number of voxels across all resolutions could indicate that ≈700 input voxels (size of the ROI at 2 mm) represents the optimal trade-off between the number of observations and input voxels, given the noise in the data and the fixed number of observations in this study.

Moreover, the present data suggest, in line with Chaimow et al. (2011), that temporal signal-to-noise-ratio, an indicator of temporal signal stability, is a critical factor for optimal decoding accuracy (Figure 3A), whereas the magnitude of BOLD signal change was not found to be relevant. While the overall BOLD signal change amplitude at 0.8 mm (4.51%) was higher than that at 2.0 mm (3.73%), the decoding performance was superior for 2.0 mm data. In fact, 0.8 mm data showed the largest magnitude of BOLD signal change but, at the same time, showed the lowest decoding accuracy among all resolutions.

Previous studies have reported BOLD response magnitude differences for different orientations. Furmanski and Engel (2000) reported that cardinal orientations elicited higher activation changes than oblique orientations of circular gratings. Swisher et al. (2010), who used the same kind of hemifield gratings as in the present study, reported higher activation for oblique than cardinal orientations. The pattern observed in this study diverges from both previous results showing a tendency for activation to be lowest for 0° and highest for 90° orientations, with oblique orientations in between. While we do not find statistically significant evidence for a differential average response magnitude across orientations at the ROI level, this does not rule out the presence of univariate orientation-discriminating signal in a subset of the input features/voxels.

It has to be noted that the comparison of decoding performance on 0.8 mm data with other resolutions is compounded by several factors. First and most importantly, the V1 coverage at this resolution was limited for technical reasons (imposed by the requirement to keep the TR at a common 2 s interval across all resolutions). This likely leads to a general underestimation of the performance at this resolution, which affects both the analyses of the full ROI, as well as those sub-sampling a smaller number of voxels. Moreover, the small coverage, combined with the impact of any residual geometric distortions, and the additional intermediate alignment step make accurate BOLD-to-structural alignment more challenging at 0.8 mm than at any other resolution. Precise alignment is important, as the V1 ROI is initially defined on the reconstructed cortical surface. Any suboptimal alignment will therefore impact decoding accuracy at 0.8 mm more than other resolutions. Lastly, the search range for C-value SVM hyperparameter was insufficient for 0.8 mm scans, the C-value was predominantly set to the lower search range boundary (Fig. S6). The search range was determined on a pilot scan and held constant for all analyses to avoid circularities. A more suitable parameter optimization scheme could have led to different results.

### Optimal low-pass spatial filtering

Gaussian spatial LP filtering is one of the most common preprocessing steps for fMRI data analyses. However, the present findings indicate that explicit spatial LP filtering, in addition to the implicit spatial filtering due to inherent motion, and the effect of head movement correction algorithms is generally not beneficial for orientation decoding (Figure 4). Only for resolutions higher than 2 mm does additional spatial smoothing with 2-3 mm FWHM show a tendency for improved decoding accuracy. This suggests that, given a resolution, a spatial smoothness equivalent to a Gaussian kernel size of ≈2 mm FWHM is optimal. This is congruent with the observation of overall lower decoding accuracies for 3 mm scans and is in line with the prediction of optimal acquisition resolution between 2 mm and 3 mm as presented above.

Moreover, spatial down-sampling is not beneficial for orientation decoding either. As shown in Figure 5 (0 mm data points, corresponding to no Gaussian smoothing), orientation decoding on down-sampled data does not outperform the decoding on data natively recorded in the corresponding resolution (as for example, in the 2.0 mm panel of Figure 5, the 0.8 mm and 1.4 mm downsampled data performed lower than native 2.0 mm data).

### Spatial characteristics of orientation specific signals

The analysis of individual spatial frequency bands via BP filtering (Fig. 4) revealed that orientation-related signal is present in a wide range of spatial frequencies as indicated by above-chance decoding performance for nearly all tested bands. However, a drop in decoding accuracy can be observed across all resolutions for bands with a 12 mm FWHM (or larger) Gaussian kernel as the smaller kernel in the LP filter pair used for BP filtering.

Freeman et al. (2013), states that it is still an open question whether fMRI can reflect signals originating from sampling random irregularities in the fine-scale columnar architecture (spatial scale ≈1 mm). This study also suggests that given a columnar architecture in the human visual cortex (Adams et al., 2007), BOLD fMRI measurements at conventional resolution ≈2 mm might reflect a combination of fine-scale and coarse-scale (spatial scale ≈10 mm) contributions. Similarly, we can interpret the present results such that the orientation-discriminating signal picked up from these BOLD fMRI data is spatially broadband in nature, includes both high spatial frequency components, as well as large-scale biases. On one hand the highest decoding accuracy was observed at 2 mm resolution, and low pass filtered components generated above chance accuracies beyond 10 mm FWHM Gaussian smoothing (similar to Op de Beeck, 2010). These observations point to indicate that low frequency components provide orientation specific signals. On the other hand we found that for DoG BP filters Gaussian kernel sizes of 4 and 5 mm FWHM and larger, decoding performance on BP filtered data was higher than the LP filtered components at all acquisition resolutions. This result pattern is an indication that low spatial frequency fMRI components also contribute to noise with respect to orientation discrimination.

According to Freeman et al. (2013), a test for fine-scale signals (≈1 mm, according to the definition by Freeman et al.) underlying the ability to decode orientations would be a comparison between decoding accuracies after down-sampling high-resolution measurements to conventional scanning resolutions, with and without prior removal of the columnar-scale contributions. To investigate this topic, we did FFT based resampling of the BOLD fMRI data from their native resolution into all three alternative resolutions with or without prior removal of low frequency components (Fig. 5). We did not observe an increase in decoding accuracy after down-sampling data from our two highest resolutions (0.8 mm and 1.4 mm), regardless of any prior LP Gaussian filtering (except for a single case of performance increase when resampling 0.8 mm to 1.4 mm data without prior LP filtering, at a comparatively low overall accuracy level). From these findings we conclude that the orientation-related signal used for decoding is unlikely to comprise of low-frequency components alone. This conclusion is in line with Swisher et al. (2010) who also reported that “majority of orientation information in high resolution fMRI activity patterns can be found at spatial scales ranging from the size of individual columns to about a centimeter”.

Carlson (2014) identified neuronal activity patterns related to stimulus edges that mimic a radial bias as a potential source of a global signal bias. The stimuli employed in this study had clearly visible, unsmoothed edges, hence edge-related activity is a valid explanation for the observed orientation-related large-scale signals. It can be argued that the V1 ROI could be adjusted by a “safety margin” to the representation of the edge of the stimuli to reduce edge related signals. We have tested various criteria for ROI definition and sizes. We have found very little variation of the results with respect to the particular shape and size of the ROI. The reported results are based on a V1 ROI generated by retinotopic mapping that used a stimulus that was larger than our visual orientation stimulus, hence we have likely sampled voxels representing edge-related signals. In other words, our ROI should contain a maximum amount of stimulus-related information present in V1. We leave an analysis exploring aspects of the relationship of individual stimulus properties and ROI shapes with the BOLD signal and decoding to a future study.

Overall, BP filtering yielded peak performances for all resolutions (except for the 3 mm acquisition). Consistent with Alink et al. (2013), the present results suggest that a band matching a DoG BP filter consisting of a 5 mm and an 8 mm FWHM Gaussian LP filter) carries most (but not all) orientation-related signal. This band covers wavelength from about 4.5 mm to 1.6 cm (Fig. S5). The Nyquist-Shannon Sampling Theorem dictates that, in order to measure a particular signal appropriately, the sampling frequency has to be at least twice the critical frequency of that signal. Hence, a 3 mm acquisition can only sample frequencies with a wavelengths of 6 mm or larger, and consequently misses some part of this most informative band.

This is consistent with our finding that optimal decoding accuracy required a resolution higher than 3 mm. The nearly identical peak performance on 1.4 mm and 2 mm data is also compatible with this minimum frequency rule. However, the markedly lower decoding performance on 0.8 mm could be considered evidence that a minimum sampling resolution is necessary but not sufficient for optimal decoding performance. In this study, an optimal balance of scanning resolution and temporal signal-to-noise-ratio is reached at 2 mm resolution. Higher resolution reduce tSNR and lower resolutions do not provide sufficient sampling of higher frequency signals.

Within the limits of our analyses the presented results do not show evidence for a variation of informative spatial frequency bands across acquisition resolutions as one might observe when a high spatial frequency signal of orientation columns in early visual cortex is reflected in (much larger) fMRI voxels by means of spatial aliasing. In the case of spatial frequency aliasing (Nyquist-Shannon Theorem) the frequency of the observed, aliased signal would vary depending on the actual sampling frequency (size of the voxel), due to an insufficient sampling frequency by the voxel grid. Here, the peak decoding performance (as found after BP filtering) is always located in the same band across all four resolutions. However, the DoG filters used here to investigate the importance of particular frequency bands feature a relatively large passband (Fig. S5) that does not allow to rule out spatial aliasing of orientation-discrimination signal. The absence of evidence for spatial aliasing is in line with Kamitani and Tong (2005) and Chaimow et al. (2011) which show that the spatial frequencies of columnar structures (0.5 cycles/mm) do not contribute signal for decoding, due to several technical limitations like inherent head motion and reduced SNR proportional to reduction in voxel volume. Moreover, Shmuel et al. (2007) state that the PSF — that captures blurring factors due to eye movements, neuronal response, BOLD response PSF in gray matter, as well as the PSF of the data acquisition process — makes fMRI data inherently LP filtered and, as such, poses a physical limitation on the spatial frequency scale from which fMRI signal can be obtained. Kamitani and Tong (2005) and Chaimow et al. (2011) identify contributions from random variations and irregularities in the columnar structures captured by larger voxels as the main source of information for decoding. These are of considerably lower frequency than the primary spatial frequency characteristics of the columnar organization and are lower than the Nyquist criterion of the BOLD fMRI sampling frequencies.

It could be speculated that the spatial scale of the orientation signal as estimated by volumetric spatial filtering is, to some degree, determined by the representation of the cortical folding pattern in the scan volume. As volumetric filtering procedures using 3D Gaussian kernels inherently mixes signals from gray matter, white matter, and superficial vessels. It might be that a volumetric BP filter corresponding to the most informative spatial frequency band is beneficial because it is of sufficient size to average signal across the entire diameter of the folded calcarine sulcus, whereas a smaller filter is not, and a bigger filter includes a substantial fraction of the surrounding white matter and adjacent cortical fields. If the above speculation is correct, we could expect lower decoding accuracy in the most informative band when replacing the employed spatial filtering procedure with a cortical surface-based smoothing or a spatial filtering that is restricted to V1 ROIs in each hemisphere. We performed these two alternative analyses and found only minor differences in the results (see supplementary material Fig. S3). Similar to the report of Swisher et al. (2010), the band-pass, high-pass, lowpass components based on these alternative spatial smoothing schemes perform very similar, but are more evenly sloped with increasing filter size compared to unconstrained volumetric filtering. Except for the 0.8 mm data, where the insufficient signal is even more evident, the BP performance is extremely similar. We conclude that there is little evidence for an impact of standard, unmasked, volumetric spatial filtering for this type of decoding analysis, compared to alternative procedures.

### Veins contribute signal usable for orientation decoding

Several studies have cited an orientation-related BOLD signal originating from the vascular system (draining veins) as a potential information source for decoding that may introduce spatial biases in the representation of orientation as measured with fMRI (Kriegeskorte et al., 2010; Chaimow et al., 2011; Shmuel et al., 2010). The present results confirm the availability of such a signal. Particularly for the two highest resolutions tested here the decoding accuracy obtained from voxels sampling veins is equal to the performance obtained from the non-venous rest of the V1 ROI, or even outperforms it when controlling for the number of input voxels for the classification model (Fig. 6A).

A BOLD signal originating in the blood vessels has the potential to introduce complex transformations of the spatial representation of orientation in the BOLD response patterns. Due to the structural properties of the vascular system this signal is likely to be of lower spatial frequency, compared to the underlying neuronal activation pattern, and is superimposed on a potential high-frequency pattern reflecting the columnar structure of V1. This explanation has been put forth by Kriegeskorte et al. (2010) who describe voxels as “complex spatio-temporal filters” and our results are compatible with this model. However, gradient echo BOLD fMRI is highly sensitive to large draining veins (Gardner, 2010; Shmuel et al., 2010; Chaimow et al., 2011), which might influence the BOLD signal also at a considerable distance from the blood vessel, rendering the interpretation of these findings even more difficult.

It should also be mentioned that previous studies found a substantial reduction of intra-vascular BOLD signals at higher magnetic field strength (Yacoub et al., 2001), and enhanced signal contributions from microvascular structures at 7T (Shmuel et al., 2007). Consequently, the particular composition of the compound signal captured with BOLD fMRI will vary with the magnetic field strength. A future study should compare the present results with data acquisitions at a different field strength to shed more light on nature of the underlying signal and the implications for decoding analysis.

#### Limitations

The focus of the present study was to investigate the effect of acquisition resolution and spatial filtering on the decoding of visual orientations from primary visual cortex. In order to yield comparable results, the acquisition parameters were constrained to guarantee a certain minimum coverage of the V1 ROI even at the highest resolutions and to have an identical temporal sampling frequency (TR) to yield the same number of observations across all resolutions. This choice implied that the GRAPPA acceleration factor had to be increased with increasing resolution, hence leading to an increased under-sampling of the k-space with higher resolutions. This could impact the sensitivity of the scan to high-frequency spatial signals. A future study will have to test whether the present findings hold when constraints on coverage and sampling frequency are relaxed. For example, a study by De Martino et al. (2013) using a 3D gradient and spin echo (GRASE) sequence suggests that such a sequence outperforms a gradient echo sequence, such as the one employed in this study, for high-resolution imaging at 0.8 mm isotropic resolution — at the expense of a vastly reduced scan volume.

The present study is exclusively based on 7 Tesla fMRI data, hence it remains unclear in which way the characteristics of the relation of decoding performance and acquisition resolution are dependent on MR field-strength. The differences in the sizes of the BOLD point-spread functions (Shmuel et al., 2007; Engel et al., 1997) suggest a lower resolution limit for 3 Tesla scans. However, the reported optimal resolution is within the range of conventional acquisition resolutions of today’s 3 Tesla scanners. A future study should address the question of how the decoding performance varies with field-strength for identical resolutions.

While this study focused on the optimal acquisition parameters for decoding of visual orientation from fMRI BOLD response patterns in early visual cortex, we acknowledge other possibilities of further optimization of the decoding procedure (classification algorithm, hyper-parameter optimization, etc.) and their potential impacts on results and interpretations. To facilitate the required future analyses we have publicly released the data (available without restrictions from GitHub https://github.com/psychoinformatics-de/studyforrest-data-multires7t) and a “Data in brief” manuscript along with this. In this study we have found that given a neural signal with known fine-scale spatial characteristics, there are technical and physiological factors that place the acquisition resolution optimal for decoding at a substantially coarser scale. Future studies should investigate whether the optimal settings for other decoding paradigms and different cortical areas, beyond the findings for visual orientation in visual cortex presented here, and the congruent results for auditory representations reported by Gardumi et al. (2016), are similar in nature.

## Acknowledgements

We thank Yukiyasu Kamitani, as well as Nikolaus Kriegeskorte and two additional anonymous reviewers their critical feedback on this research, Florian Baumgartner for his advice on data analysis, and Alex Waite for his tireless support of the computational infrastructure required for this research, and for his feedback regarding the language of this manuscript. This research was supported by a grant from the German Research Foundation (DFG) awarded to S. Pollmann and O. Speck (DFG PO 548/15-1). This research was, in part, also supported by the German Federal Ministry of Education and Research (BMBF) as part of a US-German collaboration in computational neuroscience (CRCNS; awarded to J. V. Haxby, P. Ramadge, and M. Hanke), co-funded by the BMBF and the US National Science Foundation (BMBF 01GQ1112; NSF 1129855). M. Hanke was supported by funds from the German federal state of Saxony-Anhalt and the European Regional Development Fund (ERDF), project: Center for Behavioral Brain Sciences.

## Supplementary materials and methods

### Experimental Design

This section describes how the display sequence of the oriented gratings in both the hemifields were generated per experimental run. Independent sequences were generated per hemifield with equal number of occurrences of each orientation. There were 4 different orientations (0°, 45°, 90°, or 135°) each occurring for 5 times in the sequence, contributing to 20 trials in one run. The sequences were randomly shuffled per hemifield. In this analysis a single GLM was used to model the events in both the hemifields. This was done to account for potential inter-hemispheric cross-talk due to the simultaneous bilateral stimulation, and correlation in this stimulus sequence between hemifields. Moreover, in order to minimize undesired attention shift effects, we opted for a simultaneous onsets of the stimulation in both hemifields. Combined with the further constraint of the same number of stimulation trials per orientation in both hemifields, this would unavoidably lead to a singularity of the GLM design matrix, unless a further source of temporal variability is introduced. In order to address this issue unilateral stimulation events (termed NULL events) were introduced and included in the GLM.

For comparison, we additionally analyzed these data using two separate models for both hemifields, while excluding NULL events from the modeling. This resulted in an overall improved classification performance, but did not impact the structure of the relative performance differences between resolutions (0.8 mm: 32.32%, 1.4 mm:41.78%, mm: 46.42%, and 3.0 mm: 40.17%) Figure S1 illustrates the combined impact of potential interhemispheric cross-talk and random correlations of the stimulus sequence between hemispheres by comparing the decoding performance in the contralateral and ipsilateral V1 ROI.

### Alternative spatial filtering procedures

In Figure 4 the performance of orientation decoding was quantified following lowpass, high-pass, band-pass, and band-stop spatial filtering in order to study the spatial frequency dependent orientation selective responses. All spatial filtering procedures were volumetric, using 3D Gaussian kernels and ROI voxel selection was performed after spatial filtering with different Gaussian kernel widths on the entire volume. Though this 3D filtering procedure was being extensively used in previous studies like (Op de Beeck, 2010; Swisher et al., 2010), this approach leads to information propagation from adjacent parts of the cortex, white matter and superficial vessels. Moreover, unconstrained 3D filtering does not respect the cortical folding pattern and, given a large enough filter, can smooth across sulcal boundaries, such as the two banks of the calcarine sulcus. This confounds filter width with the extent of the cortical region from which information is drawn. To avoid this problem, two additional spatial filtering approaches were implemented, namely volumetric filtering restricted to the V1 ROI, and surface-based smoothing.

#### Volumetric filtering restricted to the V1 ROI

Similar to the spatial filtering procedure performed in Alink et al. (2013), the voxel values outside the V1 ROI were considered to be missing values (NaN) instead of applying spatial filtering on the whole volume, prior to any masking. To eliminate a potential effect of smoothing across hemispheres with large Gaussian kernels, filtering was restricted to individual hemispheres. First, voxel values outside the left V1 ROI was considered to be NaNs and spatial smoothing was applied. The same procedure was applied to the right ROI, and then the smoothed left and right V1 ROI were combined to form the smoothed BOLD volume. The same nested cross validation approach was performed on the smoothed data. The results of this analysis are highly similar to the results for the unconstrained filtering prior masking (Fig. S3 A-D).

**Figure S1:**
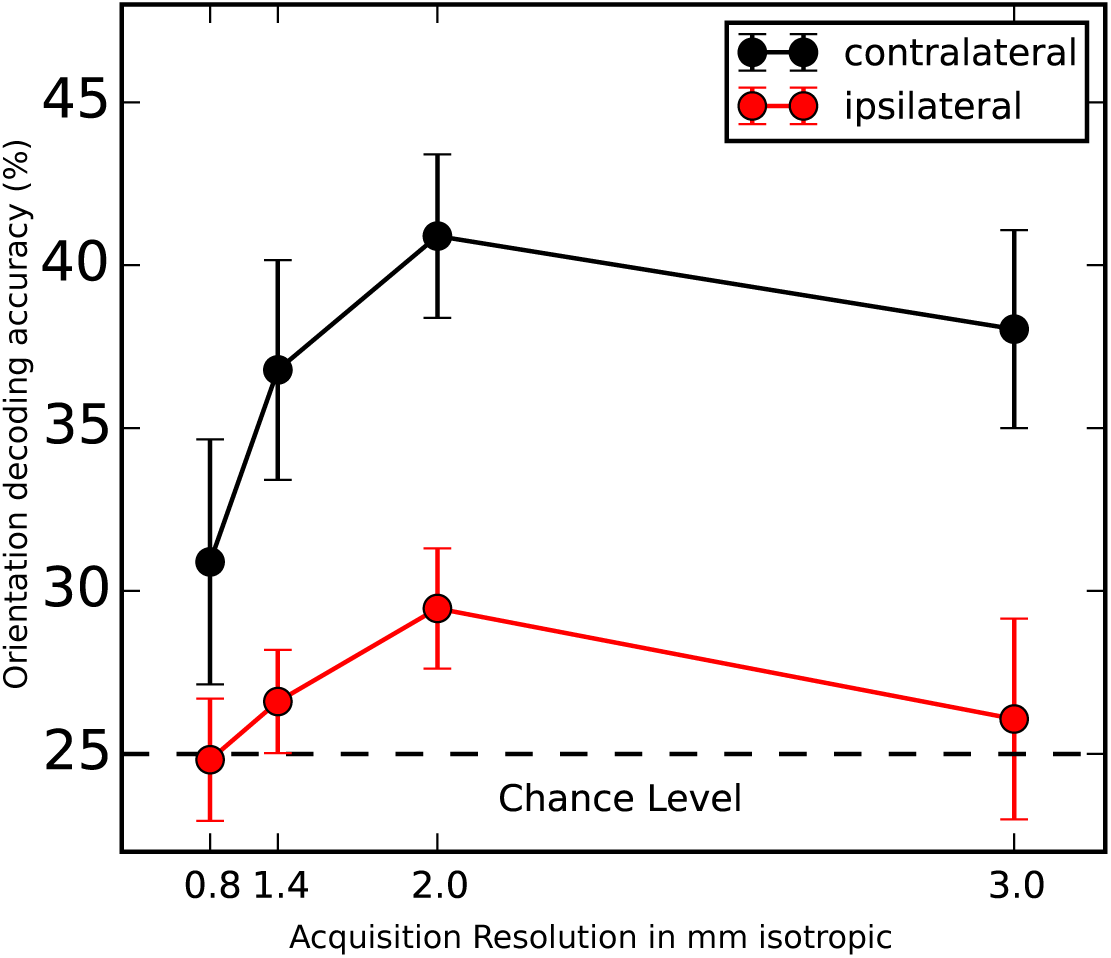
Orientation decoding accuracy on spatially unfiltered data across acquisition resolutions in both contralateral and ipsilateral V1 ROI. The ipsilateral accuracies show similar trend as the contralateral accuracies. The ipsilateral accuracies for 1.4 mm and 2 mm resolution show low decoding performance and the 0.8 mm and 3 mm decoding accuracies are at chance level.

#### Surface-based smoothing

Freesurfer’s mri vol2surf function (Dale et al., 1999) was used for smoothing gray matter BOLD data on the cortical surface, while specifying the filter size with the surf-fwhm parameter. In the next step surface-projected data were mapped back into the BOLD volume using Freesurfer’s mri surf2vol function (tri-linear interpolation, fill-projfrac parameter with range 0-1 in steps of 0.01). This procedure was performed for each hemisphere separately. Back projection into the volume was performed to maintain an equal number of input features for the decoding analysis. To illustrate the effect of surface based filtering, Figure S2 shows the reconstructed surface of one participant, with the average modeled response to cardinal and oblique orientations, filtered with 3 different filter FWHMs.

Subsequently, the same nested cross validation approach was performed on the smoothed data. The results of this analysis are shown in Figure S3 E-H. The results of surface based smoothing were similar to those of the 3D Gaussian filter, but the decoding accuracy did not decrease as rapidly with larger kernels. The band pass filtering peak was present at ≈5-8 mm but less pronounced and more evenly sloped than what was obtained from volumetric filtering.

**Figure S2:**
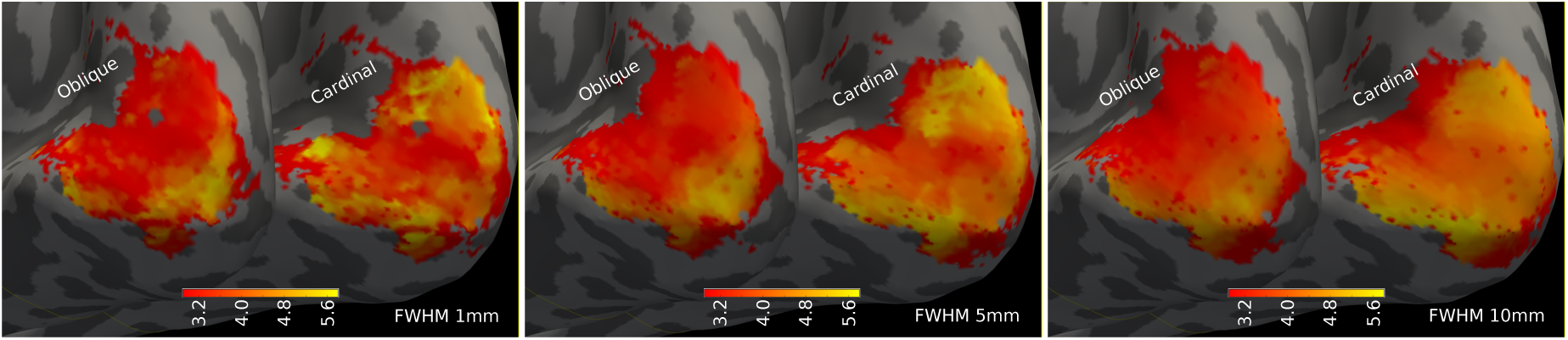
Surface-rendering of BOLD response patterns for one participant (2 mm acquisition, sub-21), after surface-based smoothing with three kernel sizes. Overlays indicate the modeled average response to cardinal and oblique orientations (*Z*-score) in the manually delineated V1 region, thresholded at *p <* 0.05 (voxelwise).

### Resampling procedure to other resolutions

Resampling BOLD fMRI data from one resolution to the other was implemented as a two-step procedure. In the following, we describe the procedure using resampling from 0.8 mm to 3.0 mm resolution as an example, but the procedure was analogous for all resolution pairs.

First FFT-based spatial filtering was performed on the distortion corrected 0.8 mm data (see Figure S4A) using the scipy function signal.resample(). This removed the higher frequency components, but the voxel grid remained unchanged (in-plane matrix size (208, 160) with 32 slices). In the next step, linear resampling/reslicing was performed with nilearn function resample img() to convert the FFT filtered image to the corresponding 3.0 mm voxel grid (see Fig. S4B for an example). Importantly, other than changing the voxel size, no further transformation, for example, to align a resampled image to the orientation of the corresponding native acquisition, were applied.

**Figure S3:**
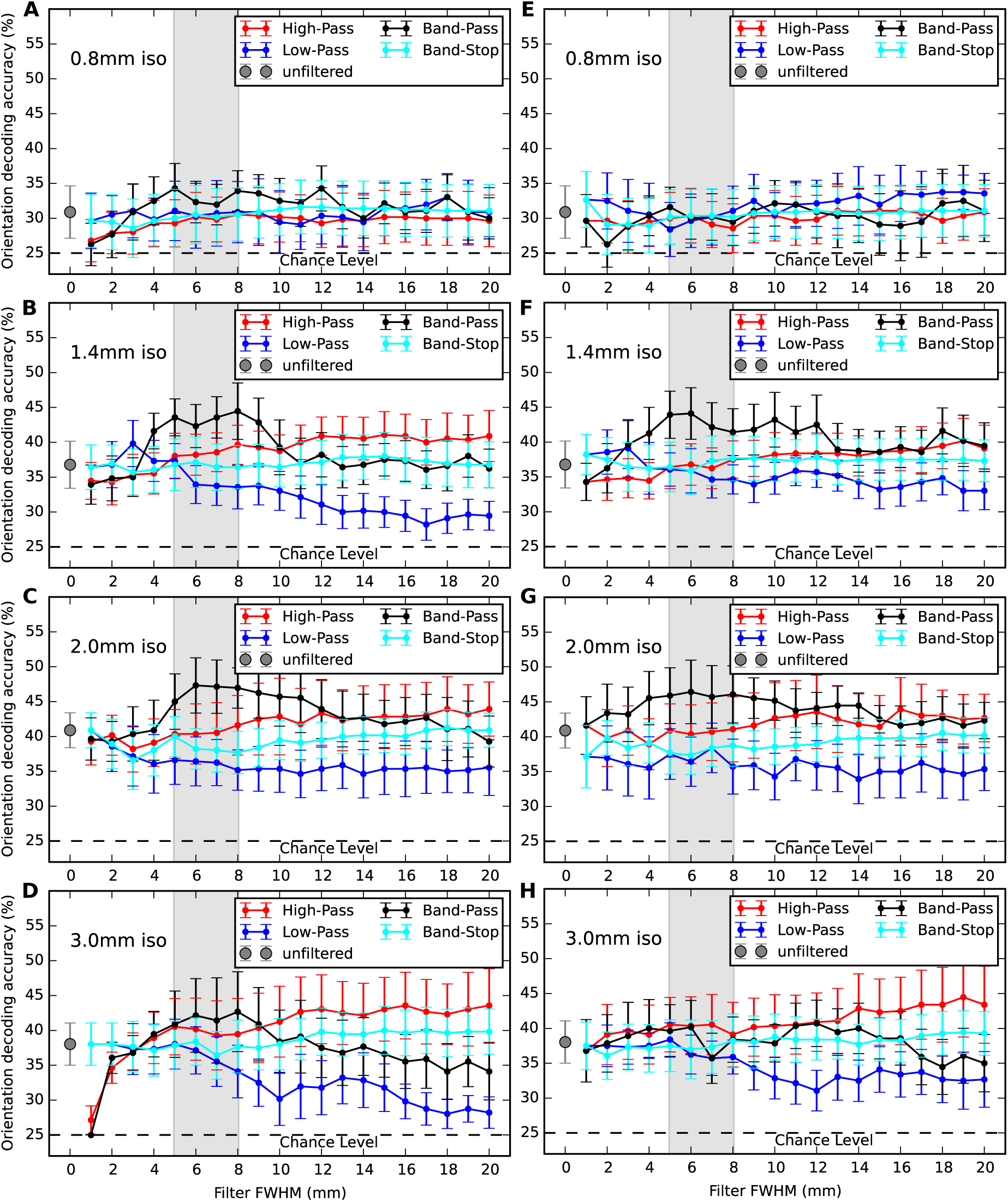
Results of alternative spatial filtering procedures (analog to Fig. 4). Volumetric spatial filtering restricted to V1 ROI (A-D), cortical surface-based smoothing (E-H).

**Figure S4:**
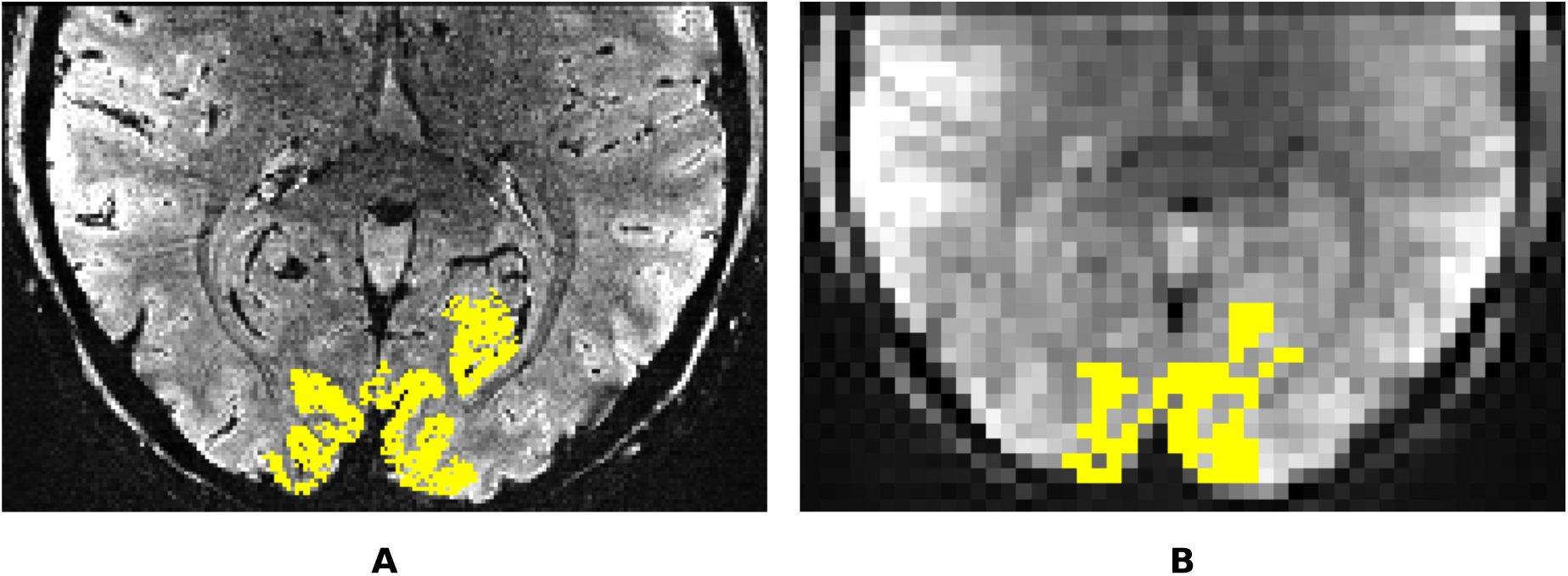
Illustration of resampling from 0.8 mm to 3.0 mm resolution. (A) Distortion corrected 0.8mm isotropic BOLD image with superimposed V1 ROI mask. (B) After removal of high-frequency components using scipy function signal.resample() superimposed with resampled V1 ROI mask (linear interpolation using scipy function ndimage.interpolation.zoom())

**Figure S5:**
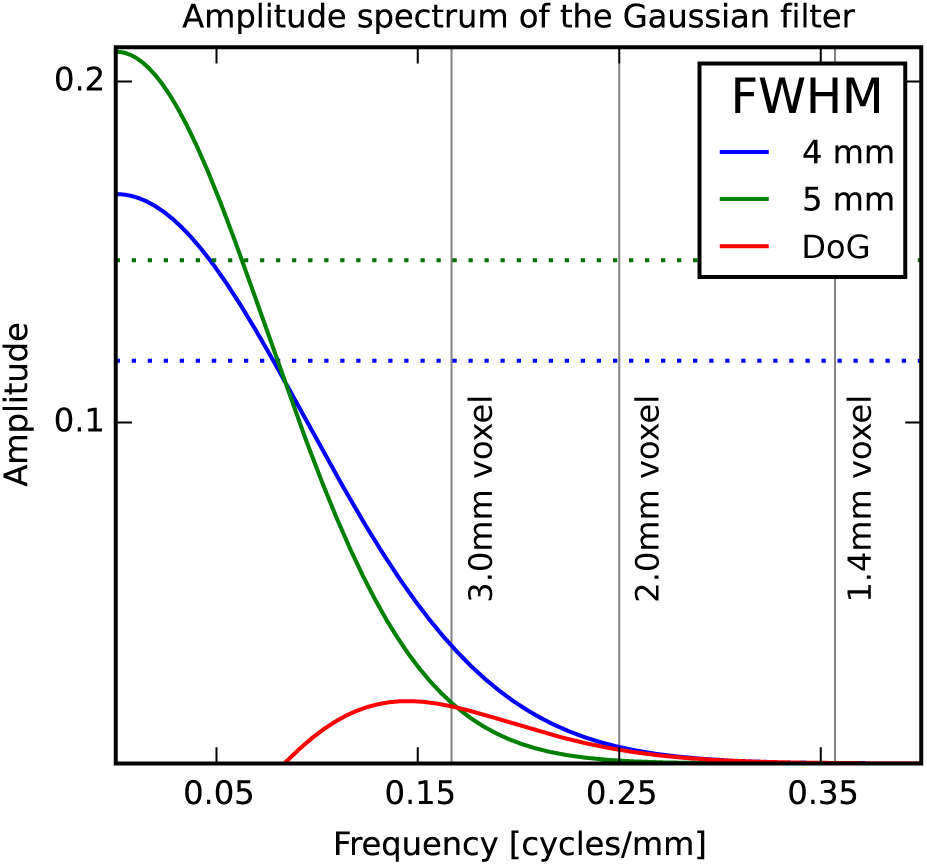
Illustration of the attenuation profile of a Difference-of-Gaussian (DoG) band-pass filter. The blue and green curve represent the profiles of Gaussian low-pass filters (4 mm and 5 mm respectively) in the frequency domain. Horizontal lines represent the −3 db points of the Gaussians. Band-pass filtering is implemented by subtracting the two low-pass filter outputs from each other. The profile of the resulting DoG band-pass filter is shown in red. Vertical lines show the Nyquist-frequencies for the three lowest resolutions in the study. The pass-band of this exemplary DoG filter (corresponding to an axis label “5 mm” in Figure 4 contains frequencies higher than what can be adequately measured with a 3 mm acquisition.

**Figure S6:**
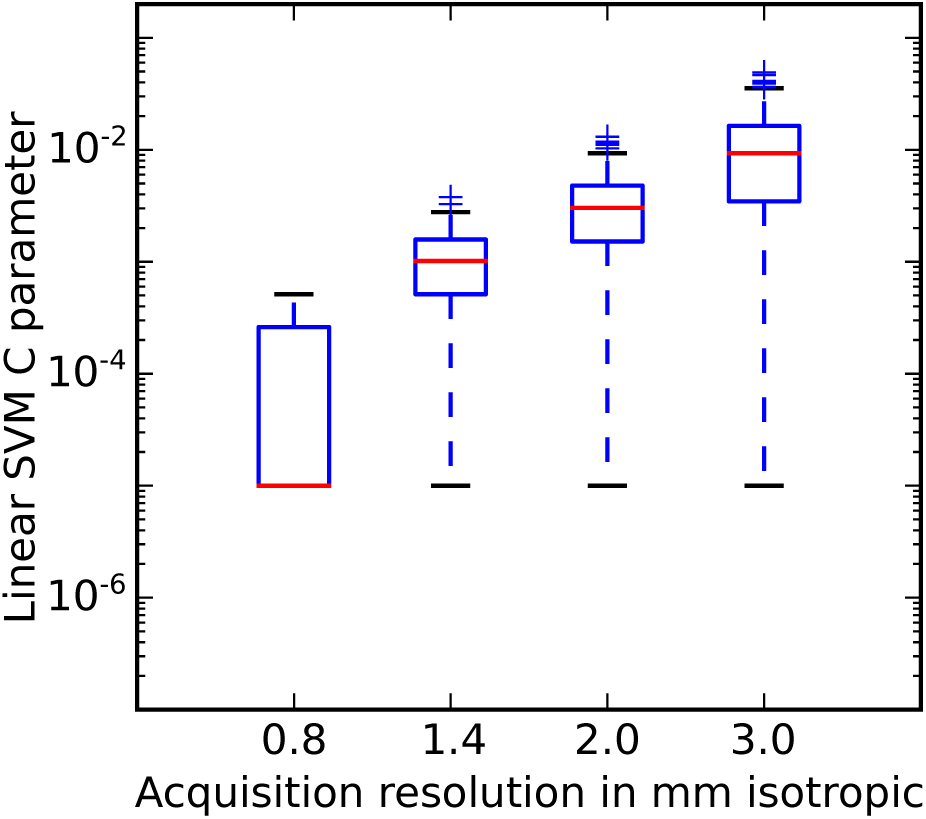
Range of tuned Linear SVM C parameters in the orientation decoding analysis across different resolutions.

**Figure S7:**
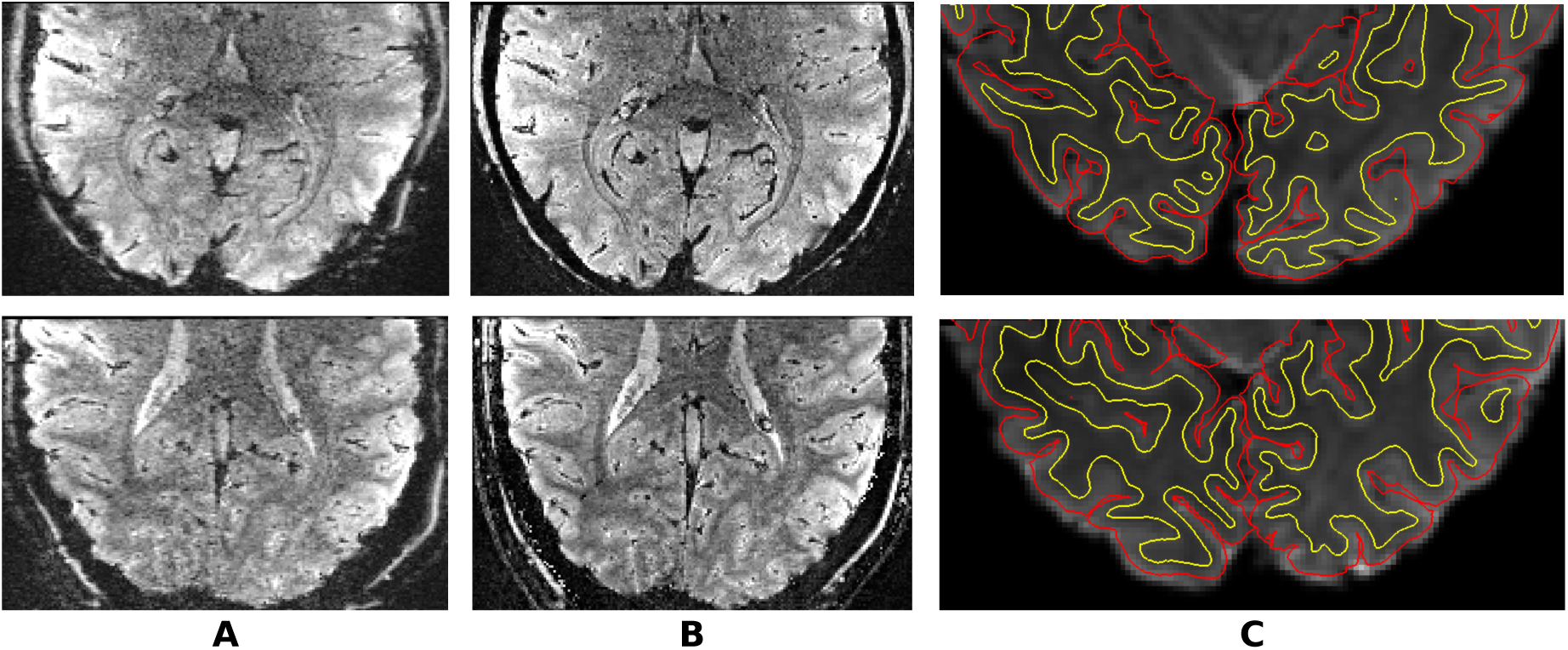
Illustration of the alignment of distortion corrected BOLD images obtained at 7 Tesla to the structural data obtained at 3 Tesla for 2 subjects. (A) Uncorrected image from Siemens 7T Magnetom (B) Distortion-corrected image (In and Speck, 2012) (C) Alignment of the BOLD image and the cortical surface, reconstructed from the corresponding structural scans. The white matter segmentation is shown in yellow and the pial surface in red.

**Figure S8:**
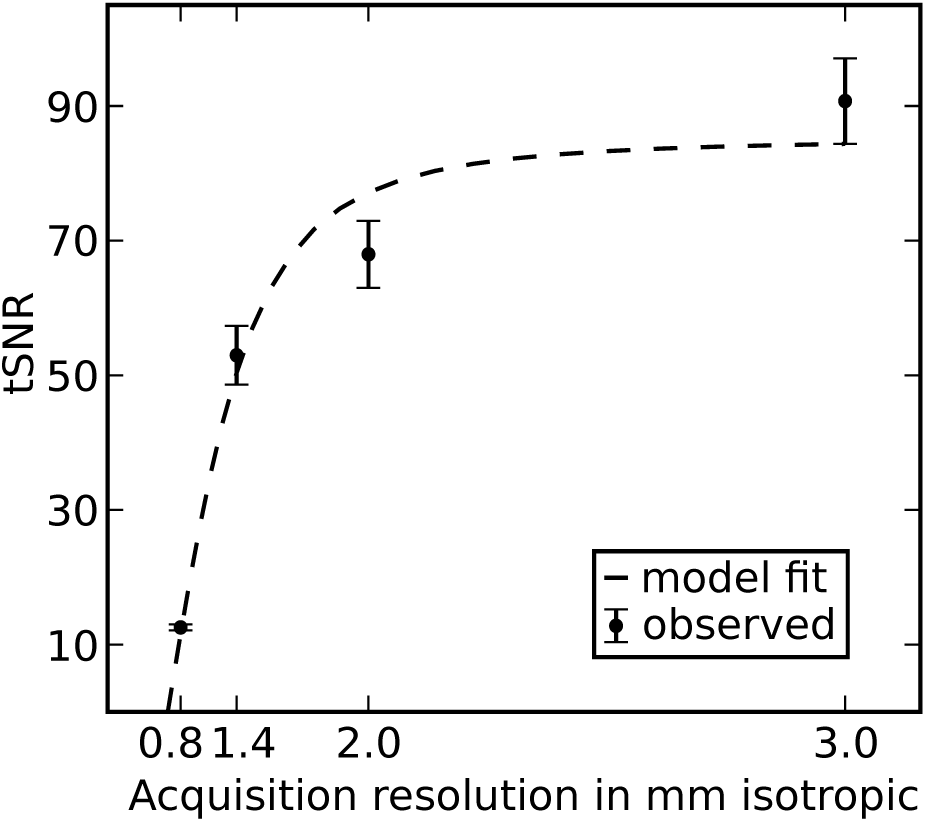
Temporal signal-to-noise ratio (tSNR) as a function of voxel volume. The observed data are represented by dots and the error bars represent the SEM across subjects. The dashed line shows the fit to the following model 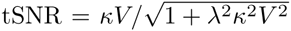 similar to the report of Triantafyllou et al. (2005)

## References

Adams, D.L., Sincich, L.C., Horton, J.C., 2007. Complete pattern of ocular dominance columns in human primary visual cortex. The Journal of Neuroscience 27, 10391–10403.

Alink, A., Krugliak, A., Walther, A., Kriegeskorte, N., 2013. fMRI orientation decoding in V1 does not require global maps or globally coherent orientation stimuli. Front Psychol 4, 493. doi:10.3389/fpsyg.2013.00493.

Op de Beeck, H., 2010. Probing the mysterious underpinnings of multi-voxel fMRI analyses. Neuroimage 50, 567–71. doi:10.1016/j.neuroimage.2009.12.072.

Bellman, R.E., 1961. Adaptive control processes: a guided tour. volume 4. Princeton University Press: Princeton.

Bonte, M., Hausfeld, L., Scharke, W., Valente, G., Formisano, E., 2014. Task-dependent decoding of speaker and vowel identity from auditory cortical response patterns. The Journal of Neuroscience 34, 4548–4557.

Carlson, T.A., 2014. Orientation decoding in human visual cortex: new insights from an unbiased perspective. The Journal of Neuroscience 34, 8373–8383. doi:10.1523/ JNEUROSCI.0548-14.2014.

Chaimow, D., Yacoub, E., Ugurbil, K., Shmuel, A., 2011. Modeling and analysis of mechanisms underlying fMRI-based decoding of information conveyed in cortical columns. Neuroimage 56, 627–42. doi:10.1016/j.neuroimage.2010.09.037.

Chang, C.C., Lin, C.J., 2011. LIBSVM: A library for support vector machines. ACM Transactions on Intelligent Systems and Technology 2, 27:1–27:27. doi:10.1145/ 1961189.1961199. Software available at http://www.csie.ntu.edu.tw/~cjlin/libsvm.

Cox, R.W., 1996. AFNI: software for analysis and visualization of functional magnetic resonance neuroimages. Computers and Biomedical research 29, 162–173. doi:10.1006/cbmr.1996.0014.

Dale, A.M., Fischl, B., Sereno, M.I., 1999. Cortical surface-based analysis: I. Segmentation and surface reconstruction. Neuroimage 9, 179–194. doi:10.1006/nimg.1998.0395.

De Martino, F., Zimmermann, J., Muckli, L., Ugurbil, K., Yacoub, E., Goebel, R., 2013. Cortical depth dependent functional responses in humans at 7t: improved specificity with 3d grase. PloS one 8, e60514. doi:10.1371/journal.pone.0060514.

Dou, W., Speck, O., Benner, T., Kaufmann, J., Li, M., Zhong, K., Walter, M., 2014. Automatic voxel positioning for MRS at 7 t. Magnetic Resonance Materials in Physics, Biology and Medicine 28, 259–270. doi:10.1007/s10334-014-0469-9.

Edwards, A.L., 1948. Note on the “correction for continuity” in testing the significance of the difference between correlated proportions. Psychometrika 13, 185–187.

Engel, S.A., Glover, G.H., Wandell, B.A., 1997. Retinotopic organization in human visual cortex and the spatial precision of functional MRI. Cerebral cortex 7, 181–192. doi:10.1093/cercor/7.2.181.

Freeman, J., Brouwer, G., Heeger, D., Merriam, E., 2011. Orientation decoding depends on maps, not columns. J Neurosci 31, 4792–804. doi:10.1523/JNEUROSCI.5160-10.2011.

Freeman, J., Heeger, D., Merriam, E., 2013. Coarse-scale biases for spirals and orientation in human visual cortex. J Neurosci 33, 19695–703. doi:10.1523/JNEUROSCI.0889-13.2013.

Friedman, J., Hastie, T., Tibshirani, R., 2001. The elements of statistical learning. volume 1. Springer series in statistics Springer, Berlin.

Furmanski, C.S., Engel, S.A., 2000. An oblique effect in human primary visual cortex. Nature Neuroscience 3, 535–536. doi:10.1038/75702.

Gardner, J.L., 2010. Is cortical vasculature functionally organized? NeuroImage 49, 1953–1956. doi:10.1016/j.neuroimage.2009.07.004.

Gardumi, A., Ivanov, D., Hausfeld, L., Valente, G., Formisano, E., Uludağ, K., 2016. The effect of spatial resolution on decoding accuracy in fmri multivariate pattern analysis. NeuroImage 132, 32–42.

Glover, G.H., 1999. Deconvolution of impulse response in event-related BOLD fMRI. Neuroimage 9, 416–429. doi:10.1006/nimg.1998.0419.

Haacke, E.M., Xu, Y., Cheng, Y.C.N., Reichenbach, J.R., 2004. Susceptibility weighted imaging (swi). Magnetic Resonance in Medicine 52, 612–618. doi:10.1002/mrm.20198.

Halchenko, Y.O., Hanke, M., 2012. Open is not enough. Let’s take the next step: An integrated, community-driven computing platform for neuroscience. Front. Neuroinform. 6. doi:10.3389/fninf.2012.00022.

Hanke, M., Baumgartner, F.J., Ibe, P., Kaule, F.R., Pollmann, S., Speck, O., Zinke, W., Stadler, J., 2014. A high-resolution 7-Tesla fMRI dataset from complex natural stimulation with an audio movie. Scientific Data 1. doi:10.1038/sdata.2014.3.

Hanke, M., Halchenko, Y., Sederberg, P., Olivetti, E., Fründ, I., Rieger, J., Herrmann, C., Haxby, J., Hanson, S., Pollmann, S., 2009. PyMVPA: A Unifying Approach to the Analysis of Neuroscientific Data. Front Neuroinform 3, 3. doi:10.3389/neuro.11.003.2009.

Haxby, J.V., 2012. Multivariate pattern analysis of fMRI: the early beginnings. Neuroimage 62, 852–855. doi:10.1016/j.neuroimage.2012.03.016.

Haynes, J., Rees, G., 2005. Predicting the orientation of invisible stimuli from activity in human primary visual cortex. Nat Neurosci 8, 686–91. doi:10.1038/nn1445.

Haynes, J.D., 2009. Decoding visual consciousness from human brain signals. Trends in cognitive sciences 13, 194–202.

Heidemann, R.M., Ivanov, D., Trampel, R., Fasano, F., Meyer, H., Pfeuffer, J., Turner, R., 2012. Isotropic submillimeter fMRI in the human brain at 7T: Combining reduced field-of-view imaging and partially parallel acquisitions. Magnetic Resonance in Medicine 68, 1506–1516. doi:10.1002/mrm.24156.

In, M.H., Speck, O., 2012. Highly accelerated PSF-mapping for EPI distortion correction with improved fidelity. Magnetic Resonance Materials in Physics, Biology and Medicine 25, 183–192.

Jenkinson, M., 2003. Fast, automated, N-dimensional phase-unwrapping algorithm. Magnetic resonance in medicine 49, 193–197. doi:10.1002/mrm.10354.

Jones, E., Oliphant, T., Peterson, P., et al., 2001. SciPy: Open source scientific tools for Python. URL: http://www.scipy.org/. [Online; accessed 2015-07-28].

Kamitani, Y., Sawahata, Y., 2010. Spatial smoothing hurts localization but not information: pitfalls for brain mappers. Neuroimage 49, 1949–52. doi:10.1016/j.neuroimage.2009.06.040.

Kamitani, Y., Tong, F., 2005. Decoding the visual and subjective contents of the human brain. Nat Neurosci 8, 679–85. doi:10.1038/nn1444.

Kriegeskorte, N., Bandettini, P., 2007. Analyzing for information, not activation, to exploit high-resolution fMRI. Neuroimage 38, 649–62. doi:10.1016/j.neuroimage.2007.02.022.

Kriegeskorte, N., Cusack, R., Bandettini, P., 2010. How does an fMRI voxel sample the neuronal activity pattern: compact-kernel or complex spatiotemporal filter? Neuroimage 49, 1965–76. doi:10.1016/j.neuroimage.2009.09.059.

Millman, K.J., Brett, M., 2007. Analysis of functional magnetic resonance imaging in Python. Computing in Science & Engineering 9, 52–55. doi:10.1109/MCSE.2007.46.

Misaki, M., Luh, W., Bandettini, P., 2013. The effect of spatial smoothing on fMRI decoding of columnar-level organization with linear support vector machine. J Neurosci Methods 212, 355–61. doi:10.1016/j.jneumeth.2012.11.004.

Obermayer, K., Blasdel, G.G., 1993. Geometry of orientation and ocular dominance columns in monkey striate cortex. The Journal of Neuroscience 13, 4114–4129.

Pedregosa, F., Varoquaux, G., Gramfort, A., Michel, V., Thirion, B., Grisel, O., Blondel, M., Prettenhofer, P., Weiss, R., Dubourg, V., et al., 2011. Scikit-learn: Machine learning in Python. The Journal of Machine Learning Research 12, 2825–2830.

Peirce, J.W., 2008. Generating stimuli for neuroscience using PsychoPy. Frontiers in neuroinformatics 2. doi:10.3389/neuro.11.010.2008.

Pereira, F., Mitchell, T., Botvinick, M., 2009. Machine learning classifiers and fMRI: a tutorial overview. Neuroimage 45, S199–S209. doi:10.1016/j.neuroimage.2008.11.007.

Sengupta, A., Kaule, F., Guntupalli, J.S., HHoffmann, M., Häusler, C., Stadler, J., Hanke, M., 2016. An extension of the studyforrest dataset for vision research. Scientific Data, Under reviewURL: http://biorxiv.org/content/early/2016/03/31/046573.

Shmuel, A., Chaimow, D., Raddatz, G., Ugurbil, K., Yacoub, E., 2010. Mechanisms underlying decoding at 7T: ocular dominance columns, broad structures, and macroscopic blood vessels in V1 convey information on the stimulated eye. Neuroimage 49, 1957–1964. doi:10.1016/j.neuroimage.2009.08.040.

Shmuel, A., Yacoub, E., Chaimow, D., Logothetis, N., Ugurbil, K., 2007. Spatiotemporal point-spread function of fMRI signal in human gray matter at 7 Tesla. Neuroimage 35, 539–52. doi:10.1016/j.neuroimage.2006.12.030.

Smith, S., Jenkinson, M., Woolrich, M., Beckmann, C., Behrens, T., Johansen-Berg, H., Bannister, P., De, L.M., Drobnjak, I., Flitney, D., Niazy, R., Saunders, J., Vickers, J., Zhang, Y., De, S.N., Brady, J., Matthews, P., 2004. Advances in functional and structural MR image analysis and implementation as FSL. Neuroimage 23 Suppl 1, S208–19. doi:10.1016/j.neuroimage.2004.07.051.

Swisher, J., Gatenby, J., Gore, J., Wolfe, B., Moon, C., Kim, S., Tong, F., 2010. Multiscale pattern analysis of orientation-selective activity in the primary visual cortex. J Neurosci 30, 325–30. doi:10.1523/JNEUROSCI.4811-09.2010.

Tong, F., Harrison, S., Dewey, J., Kamitani, Y., 2012. Relationship between BOLD amplitude and pattern classification of orientation-selective activity in the human visual cortex. Neuroimage 63, 1212–22. doi:10.1016/j.neuroimage.2012.08.005.

Triantafyllou, C., Hoge, R., Krueger, G., Wiggins, C., Potthast, A., Wiggins, G., Wald, L., 2005. Comparison of physiological noise at 1.5 T 3 T and 7 T and optimization of fMRI acquisition parameters. NeuroImage 26, 243–250. doi:10.1016/j.neuroimage.2005.01.007.

Uğurbil, K., 2012. The road to functional imaging and ultrahigh fields. Neuroimage 62, 726–35. doi:10.1016/j.neuroimage.2012.01.134.

Warnking, J., Dojat, M., Guérin-Dugué, A., Delon-Martin, C., Olympieff, S., Richard, N., Chéhikian, A., Segebarth, C., 2002. fMRI retinotopic mapping–step by step. Neuroimage 17, 1665–83. doi:10.1006/nimg.2002.1304.

Weibull, A., Gustavsson, H., Mattsson, S., Svensson, J., 2008. Investigation of spatial resolution, partial volume effects and smoothing in functional MRI using artificial 3d time series. NeuroImage 41, 346–353. URL: http://dx.doi.org/10.1016/j.neuroimage.2008.02.015, doi:10.1016/j.neuroimage.2008.02.015.

Yacoub, E., Harel, N., Ugurbil, K., 2008. High-field fMRI unveils orientation columns in humans. Proc Natl Acad Sci U S A 105, 10607–12. doi:10.1073/pnas.0804110105.

Yacoub, E., Shmuel, A., Pfeuffer, J., De Moortele, V., Adriany, G., Andersen, P., Vaughan, J.T., Merkle, H., Ugurbil, K., Hu, X., et al., 2001. Imaging brain function in humans at 7 tesla. Magnetic Resonance in Medicine 45, 588–594.

Zhang, F., Wang, J.P., Kim, J., Parrish, T., Wong, P.C., 2015. Decoding multiple sound categories in the human temporal cortex using high resolution fmri. PloS one 10, e0117303.

